# Ventricular, atrial and outflow tract heart progenitors arise from spatially and molecularly distinct regions of the primitive streak

**DOI:** 10.1101/2020.07.12.198994

**Authors:** Kenzo Ivanovitch, Pablo Soro-Barrio, Probir Chakravarty, Rebecca A Jones, Donald M Bell, S. Neda Mousavy Gharavy, Despina Stamataki, Julien Delile, James C Smith, James Briscoe

## Abstract

The heart develops from two sources of mesoderm progenitors, the first and second heart field (FHF and SHF). Using a single cell transcriptomic assay in combination with genetic lineage tracing, we find the FHF and SHF are subdivided into distinct pools of progenitors in gastrulating mouse embryos at earlier stages than previously thought. Each subpopulation has a distinct origin in the primitive streak. The first progenitors to leave the primitive streak contribute to the left ventricle, shortly after right ventricle progenitor emigrate, followed by the outflow tract and atrial progenitors. Moreover, a subset of atrial progenitors are gradually incorporated in posterior locations of the FHF. Although cells allocated to the outflow tract and atrium leave the primitive streak at a similar stage, they arise from different regions. Outflow tract cells originate from distal locations in the primitive streak while atrial progenitors are positioned more proximally. Moreover, single cell RNA sequencing demonstrates that the primitive streak cells contributing to the ventricles have a distinct molecular signature from those forming the outflow tract and atrium. We conclude that cardiac progenitors are pre-patterned within the primitive streak and this prefigures their allocation to distinct anatomical structures of the heart. Together, our data provide a new molecular and spatial map of mammalian cardiac progenitors that will support future studies of heart development, function and disease.

## Introduction

The cells that comprise different parts of an organ can arise from distinct origins and acquire their fate at different times during ontogeny. In the case of the heart, two mesodermal derived sources of cardiac progenitors, named the first heart field (FHF) and the second heart field (SHF), have been identified (reviewed in Meilhac and Buckingham, 2018). The FHF forms mainly the left ventricle and constitutes the initial cardiac crescent. By contrast, the cardiac progenitors of the SHF contribute to the right ventricle, outflow tract and atria in addition to branchiomeric muscles.

Retrospective clonal analysis has suggested that the FHF and SHF progenitors segregate either before or at the onset of gastrulation (Meilhac and Buckingham, 2018; Meilhac et al., 2004). The analysis showed large clones spanning multiple compartments. Clones belonging to the first lineage labelled the left ventricle and other compartments with the exception of the outflow tract while the second lineage contributed to the outflow tract, atria and right ventricle but never to the left ventricle. Subclones within the first and second lineages were also observed. These were smaller in size and suggested that the individualisation of the different regions of the heart happens at later stages (Meilhac et al., 2004).

Consistent with the notion that the cardiac lineages are established during gastrulation, clonal analysis based on tracing the progeny of *Mesp1*-expressing cells, which is expressed in the primitive streak and nascent mesoderm (https://marionilab.cruk.cam.ac.uk/MouseGastrulation2018/; Arnold and Robertson, 2009; Costello et al., 2011; Pijuan-Sala et al., 2019), indicated that independent sets of *Mesp1* expressing cells contribute to FHF and SHF derivatives (Devine et al., 2014; Lescroart et al., 2014). These can be distinguished by their time of appearance during gastrulation, with embryonic day (E)6.5 *Mesp1+* cells supplying the FHF while SHF derivatives preferentially derive from E7.5 *Mesp1*+ cells (Lescroart et al., 2014). The *Mesp1* clonal analysis resulted in clones of small sizes restricted to compartments derived from the FHF or SHF (Devine et al., 2014; Lescroart et al., 2014). Thus, these observations are consistent with the existence of two groups of progenitors that subsequently became further restricted to specific cardiac fates in the mesoderm.

In addition to a separation between the FHF and SHF, genetic tracing experiments uncovered a distinction between the atria and ventricular progenitors. Genetic lineage tracing with the transcription factor *Foxa2* identifies a population of cardiac progenitors in the anterior primitive streak at E6.5 that contribute to both left and right ventricles but not to the atria (Bardot et al., 2017). This is in line with fate mapping studies in the chick showing that the atrial and ventricular cells arise at different anterior-posterior positions in the primitive streak (Garcia-Martinez and Schoenwolf, 1993; Yutzey and Bader, 1995). Although these analyses did not resolve the clonal relationship between the cells, these findings are compatible with a model in which sublineages within the FHF and SHF lineages already exist in the primitive streak, such that the atrial lineage is distinct from the right ventricle lineage within SHF progenitors. This would imply that two distinct cardiac progenitors exist solely at the epiblast stage, prior to gastrulation, and that more subpopulations of cardiac progenitors than first and second progenitors can be molecularly defined in the primitive streak. The most rigorous way to decide if this is the case is to define the location of all the cardiac progenitors in the streak and the precise embryonic stages at which they ingress. This would allow the comparison of distinct subpopulations of cardiac precursors and reveal putative molecular differences of cardiac progenitors in the primitive streak. Single cell transcriptomic assays show *Mesp1+* cardiac progenitors diversify into molecularly distinct FHF and SHF populations at late embryonic day (E7.25) stage once cells have migrated and reached their final location in the embryo (Lescroart et al., 2018). The signalling environment cells encounter during and after their migration might play a role in the patterning of the cardiac progenitors into FHF and SHF domains (Andersen et al., 2018; Bardot et al 2020; Guzzetta et al., 2020; Hochgreb et al., 2003; Xavier-Neto et al., 1999). Whether initial molecular differences between different sets of cardiac progenitors already exist in the primitive streak remains unclear.

In this study, we use genetic tracing of *T* and *Foxa2* expressing cells and find there is an orderly allocation of primitive streak cells first into the left ventricle at mid-streak stage, then the right ventricle progenitors at late streak stage, and finally at the no bud (OB)-early bud (EB) stage into outflow tract and atrial progenitors. Consistent with this, we identified independent sets of *Foxa2*+ cells that are allocated to the left ventricle and right ventricle. Allocation of cells to the outflow tract and atria happens at similar gastrulation stages but from distinct locations within the primitive streak. The outflow tract originates from distal regions of the primitive streak, while proximal regions contribute to the atria. Moreover, the outflow tract forms from primitive streak cells that initially expressed *Foxa2*, but subsequently turned off *Foxa2* expression as they switched their contribution from right ventricle to the outflow tract. Crucially, by combining single cell transcriptomic assays with a lineage tracer to label cells supplying only the poles of the heart, we uncover molecularly distinct subpopulations of cells that correspond to progenitors for right ventricle, outflow tract and atria in the SHF. Further single cell transcriptomic experiments at earlier stages established that the primitive streak cells contributing to the ventricles and outflow tract/atria are also molecularly distinct. Thus, rather than a simple subdivision of cardiac progenitors into FHF and SHF, our analysis reveals a more elaborate map for the source of cells that form the heart with distinct spatial and temporal origins for outflow tract and atrial progenitors as well as left and right ventricular progenitors. We conclude that the cardiac progenitors are pre-patterned within the primitive streak and this prefigures their contribution to distinct anatomical structures of the heart both in time and space. This has implications for the classification of congenital heart diseases based on the origin of malformation in a specific mesodermal lineage as well as for the design of *in vitro* methods to generate specific cardiac cells from pluripotent stem cells.

## Results

### Genetic tracing of primitive streak cells using a tamoxifen inducible T reporter

During gastrulation, epiblast cells ingress through the primitive streak to form the mesoderm. This process occurs over an extended period of time, from embryonic day (E)6 to E8 in the mouse. The T-box transcription factor *T* is expressed in the primitive streak and is downregulated shortly after ingression during migration within the nascent mesoderm. To assess the developmental time points at which the *T*-expressing primitive streak cells are destined to contribute to the heart, we performed genetic tracing using an inducible *T^nGFP-CreERT2/+^* mouse, expressing *CreERT2* and nuclear localised GFP (nGFP) downstream of the endogenous *T* (Imuta et al., 2013), in combination with the *R26R^tdTomato/+^* reporter mice (*T^nEGP-CreERT2/+^; R26R^tdTomato/+^*, Fig 1A) (Madisen et al., 2010).

**Fig 1.**
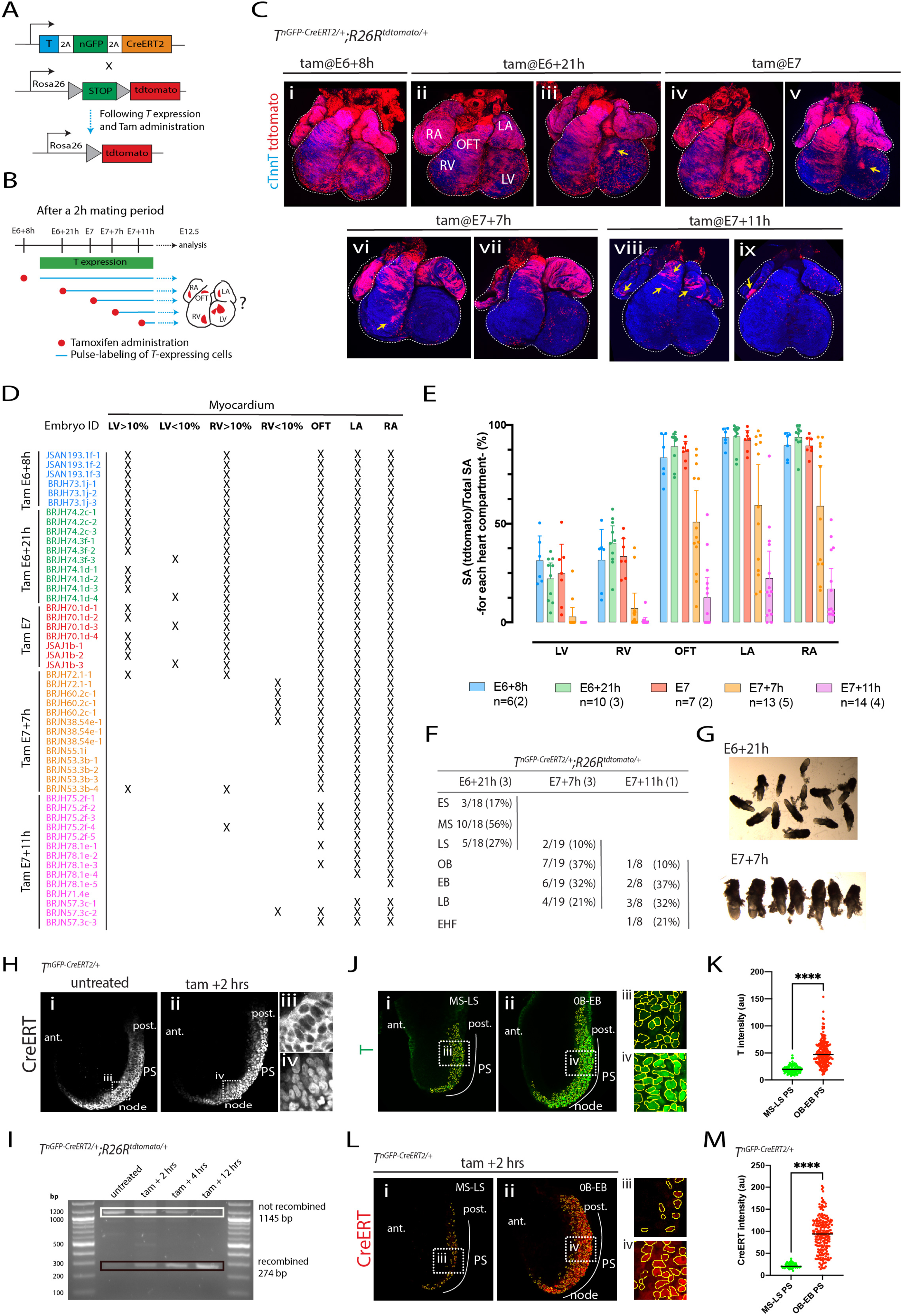
Genetic tracing of the *T+* primitive steak cells. **(A)** Schematics of the *T^nGPF-CreERT2/+^; R26R^tdTomato/+^* allele (Imuta et al., 2013). Cre-ERT2 is expressed in cells expressing *T*. In the presence of tamoxifen, Cre protein is translocated to the nucleus where it recombines the *R26R^tdTomato/+^* reporter. As a result, the cell and its descendants are permanently labeled. **(B)** Diagram of the experimental approach. *T*-expressing cells and their descendants are labelled, from E6+21h, E7, E7+7h and E7+11h by administrating a dose of tamoxifen (Tam) to *T^nGPF-CreERT2/+^; R26R^tdTomato/+^* mice (0.08mg/body weight via oral gavage). Cell descendants in the myocardium are analysed at E12.5. **(C)** Representative hearts resulting from the administration of tamoxifen at different time points in *T^nGPF-CreERT2/+^; R26R^tdTomato/+^* immunostained with cTnnT to reveal the cardiomyoyctes (blue). Yellow arrows identify small patches of tdTomato positive cardiomyocytes in the LV (iii and v), in the RV (v) and in the outflow tract and atria (vii and viii). Views are ventral. Single epicardial cells are labeled in each condition. **(D-E)** Summary of all *T^nGPF-CreERT2/+^; R26R^tdTomato/+^* hearts examined. The contribution of the *T-*expressing cells to the different compartments of the heart is quantified by measuring the proportion of tdTomato positive myocardium. Numbers in brackets in (E) represent the number of litters assessed. Error bars are SD. **(F)** Stage variation quantified according to Downs and Davies criteria (Downs and Davies, 1993). Number in brackets represents the number of litters assessed. All the timed matings were for 2-hour periods. (**G**) Embryos collected at E6+21h or E7+7h showing variation in stage. ((H) Representative *T^nGPF-CreERT2/+^*embryos untreated or tamoxifen treated for two hours mice (0.08mg/body weight via oral gavage) and immunostained with estrogen receptor. Insets iii and iv are magnified view from i and ii respectively. (**I**) PCR amplicons generated from the genomic region in which Cre-mediated recombination occurs, resolved on an agarose gel. Before recombination, the PCR product is 1145 bp (white rectangle); after recombination it is 274 bp (black rectangle). Template gDNA was extracted from either an ear clip of an adult *Tn^GPF-CreERT2/+^*; *R26^RtdTomato/tdTomato^* mouse (untreated) or *T^nGPF-CreERT2/+^*; *R26R^tdTomato/tdTomato^* embryos dissected at 2, 4 and 12 hours following oral gavage with Tamoxifen, as labelled. An increase in the proportion of the recombined band can be seen over time following Tamoxifen administration. (J) Representative *T^nGPF-CreERT2/+^*embryos at the MS-LS and OB-EB stages, immunostained with. (iii) and (iv) are magnified views in (i) and (ii) respectively. (K) Quantification of T intensity in single segmented nuclei from embryo shown in (J). (L) Representative *T^nGPF-CreERT2/+^*embryos at the MS-LS and OB-EB stages treated with tamoxifen for two hours (0.08mg/body weight via oral gavage) and immunostained for estrogen receptor. (iii) and (iv) are magnified views in (i) and (ii) respectively. (M) Quantification of Estrogen receptor intensity in single segmented nuclei from the embryos shown in (L). Embryos in J (i) and (ii) and in L (i) and (ii) were immunostained together and imaged under similar conditions. LV: left ventricle, RV: right ventricle, OFT: outflow tract, LA: left atria, RA: right atria, cTnnT: cardiac troponinin T. MS: Mid-streak LS: late-streak, OB: “no bud” stage, EB: “early bud” stage, LB: “late bud stage”, EHF: early head fold. cTnnT: cardiac Troponin T. ant.: anterior, post.: posterior, PS: primitive streak. Scale bar: 200 µm.

We first assessed how long the activity of the tamoxifen persists after it is administered by oral gavage in a pregnant mouse. We administered tamoxifen (0.08mg/bw) at E5, ∼24 hours prior to the initial onset of *T*-expression in the primitive streak (Rivera-Perez and Magnuson, 2005), and we detected recombined tdTomato-expressing cells within mesodermal derivatives including the heart tube, head mesenchyme and endothelium, albeit at a low density, in E8.5 embryos, (S1A Fig.). Thus, the activity of the tamoxifen persists for at least 24 hours at this dose.

### Primitive streak cells contribute first to the ventricles and subsequently to the outflow tract and atria myocardium

We next administered single doses of tamoxifen *T^nEGP-CreERT2/+^; R26R^tdTomato/+^* at successive stages of gastrulation. We reasoned this would result in pulse-labelling of most populations of *T*-expressing cells from the time of administration for at least 24 hours (Fig 1B). The fate of their progeny could then be followed and the developmental stages during which primitive streak cells contribute to the left and right ventricle, atria and outflow tract could be deduced. To gain better control over the embryonic stages, mice were synchronized in estrus and mated over short periods of 2 hours for all the experiments (from 7am to 9 am; vaginal plugs were checked at 9 am and if positive, were defined as E0). To quantify the contribution of the *T*-expressing cells to the heart, we measured surface area occupied by the tdTomato-positive cells in the heart myocardium at E12.5 and within each cardiac chamber.

An early administration of tamoxifen at E6+8h resulted in the population of left and right ventricles, outflow tract and atria with tdTomato-expressing cardiomyocytes (in 6 out of 6 hearts analysed, Fig 1Ci, D and E). The contribution of tdTomato-positive cardiomyocytes to the left and right ventricle was similar (left ventricle: 31.4% ±12.4; right ventricle: 31.6% ± 15.5, mean ± SD, p-value: ns). Contribution to the outflow tract and atria was the highest, covering almost their entirety in all cases (83.8% ± 11.7 and 91.7 ± 5.9, mean ± SD, for outflow tract and atria, respectively).

An administration at E6+21h or E7 also resulted in tdTomato-expressing cardiomyocytes in the left and right ventricles, outflow tract and atria (in 17 out of 17 hearts analysed, Fig 1Cii-v, D and E). In these cases, the contribution of the tdTomato-positive cells to the left ventricle was lower, on average, than to the right ventricle (for E6+21h, left ventricle: 22.2% ±11.1; right ventricle: 40.2% ± 12.2 and for E7, left ventricle: 24.8% ±16, and right ventricle: 33.4% ± 9.9, mean ± SD, p-value: 0.004). Contribution to the outflow tract and atria continued to be the highest in all cases (89.1%± 8.4 and % 94.1± 6.9, mean ± SD, for outflow tract and atria, respectively). Variability in embryonic stages within litters at the time of tamoxifen administration can explain the variability in the results (see below). For example, in 4/17 hearts, tdTomato-expressing cardiomyocytes populated less than 10% of the total left ventricle surface area. In those hearts, the proportion of tdTomato-positive cardiomyocytes found in the right ventricle was high (Fig 1Ciii, v and D). We never observed the reverse, however, where a lower proportion of tdTomato positive cardiomyocytes was found in the right ventricle compared to the left ventricle (Fig 1D). These results suggest that subsets of the embryos at E6+21h and E7 were at more advanced embryonic stages with most left ventricle precursors having already left the primitive streak with right ventricular precursors remaining in the primitive streak.

We next asked if the administration of tamoxifen at a later time point would result in the absence of tdTomato-expressing cardiomyocytes in the left ventricle and right ventricle. We administered tamoxifen at E7+7h. TdTomato-expressing cardiomyocytes were detected in the left ventricle in only 2/13 hearts (Fig 1D). TdTomato-positive cardiomyocytes were detected more frequently in the right ventricle, in 7/13 hearts (Fig 1D). However, these cells covered less than 5% of the total right ventricle surface area (yellow arrow in Fig 1Cvi and E), except for the two hearts in which tdTomato-expressing cardiomyocytes were also detected in the left ventricle. These results indicate that left ventricle and most right ventricle precursors have left the primitive streak by E7+7h. In these embryos, tdTomato-expressing cardiomyocytes were evident in the outflow tract and atria in 13/13 hearts, covering in some cases between 72.8% and 94% of the total surface area of the outflow tract and atria, including hearts without tdTomato-expressing cardiomyocytes in the left ventricle and right ventricle. These results suggest that the majority of the outflow tract and atria precursors are still located in the primitive streak at stages after the ventricular precursors have already left the primitive streak.

Finally, when tamoxifen was administered at a later stage (E7+11h), the contribution of the tdTomato-expressing cardiomyocytes to the outflow tract and atria was lower (Fig 1Cviii-ix and E, 12.6% ± 17.4 and 19.7% ± 20.8). In all cases, these embryos lacked tdTomato-expressing cells in the left and right ventricles. In 5/14 hearts, tdTomato-expressing cells were absent in the outflow tract but present in the atria albeit in low numbers. The reverse, *i.e*. tdTomato expressing cells found only in the outflow tract but not in the atria, was never observed, suggesting that the atrial precursors are the last cardiomyocyte precursors to leave the primitive streak.

Combining the *T^nEGP-CreERT2/+^* transgene with the R26^mGFP/+^ reporter (Muzumdar et al 2007) produced similar results (S2A Fig). An early tamoxifen administration (at E6+8h) led to mGFP positive cardiomyocytes populating the left ventricle myocardium. When tamoxifen was administered late (E7+7h) no mGFP cardiomyocytes were identified in the left ventricle myocardium. mGFP cardiomyocytes were observed in the outflow tract and atria in both conditions.

To analyse variability in embryonic stages at the time of tamoxifen administration, we dissected litters of *T^nEGP-CreERT2/+^; R26R^tdTomato/+^* embryos at E6+21h and E7+7h and staged embryos according to morphological landmarks using the dissecting brightfield microscope (Downs et al., 1993). As expected, we found a range of stages for each time point. Litters dissected at E6+21h included early (17%) mid (56%) and late-streak stages (27%). Litters dissected at E7+7h included late streak (10%), no bud (OB, 37%), early bud (EB, 32%) and late bud stages (LB, 21%) (Fig 1F and G). Whole mount immunostaining for the oestrogen receptor ERT revealed CreErt2 in the nucleus within two hours of oral gavage of tamoxifen (0.08mg/bw) (Fig 1H and S3Ai-vi Fig.). PCR analysis showed that the *R26R^tdTomato/+^* locus was recombined (Fig 1I and S4Ai-ii and B Fig.).

Together, these results indicate that cardiac progenitors ingress in an orderly sequence through the primitive streak. Left ventricle progenitors leave the primitive streak first, at the mid-streak stage, followed by right ventricle progenitors at the late-streak stage. Outflow tract and atrial precursors leave the primitive streak at subsequent stages, starting around the OB stage.

### *T* is expressed at low level in ventricular progenitors and at higher levels in atrial and outflow tract progenitors

Ventricular progenitors were captured less efficiently than the OFT and atria progenitors in *T*-lineage tracing experiments. This prompted us to quantify expression levels of T and Cre protein in embryos collected at the mid to late streak, when ventricular progenitors reside in the streak, and OB to EB, when OFT and atria progenitors reside instead in the streak. All primitive streak cells express *T*, however, we found that primitive streak cells at the mid to late streak stages express lower levels of T protein compared to the OB and EB stages (Fig 1Ji-iv and K). The transgenic *T^nEGP-CreERT2/+^* embryo showed a similar pattern of expression, with lowest Cre (Fig 1L and M) and nGFP signal at the mid-late streak stage (compare GFP levels in S4Ai-ii Fig). These results suggest why not all ventricular progenitors are captured by our *T*-lineage tracing experiments.

### Late primitive streak cells/atrial progenitors contribute to posterior regions of the first heart field and differentiate early into cardiomyocytes

The first heart field contributes mainly to the left ventricle in addition to a minority of the atria (Cai et al. 2003, Spater et al. 2013) and our late *T*-lineage tracing at E7+7h may have missed these FHF atrial progenitors. This prompted us to test whether the descendants from the late E7+7h were also populating the first heart field. We assayed cTnnT, to mark first heart field cells, since these differentiate early into cardiomyocytes to establish the cardiac crescent. We found in 5 out of 9 embryos analysed, sparse tdTomato-expressing cardiomyocytes located in posterior regions of the cardiac crescent or prospective inflow (Fig. 2Ai-iv). No tdTomato-expressing cardiomyocytes were found in anterior regions of the cardiac crescent in any of the embryos examined. TdTomato positive cells could also be found in the endoderm (blue arrows in Fig 2Aii). Furthermore, analysis at the heart tube stage reveals that late E7+7h T cell descendants were populating the inflow regions of the heart tube (in 3 out of 4 embryos, Fig 2Bi and yellow arrows in Fig 2Bii). A sparse contribution to the endocardium was also noted (blue arrows in Fig 2Biii). We conclude that a subset of the late *T*-expressing cells/atrial progenitors are recruited to posterior regions of the first heart field and differentiate early into cardiomyocytes.

**Fig 2.**
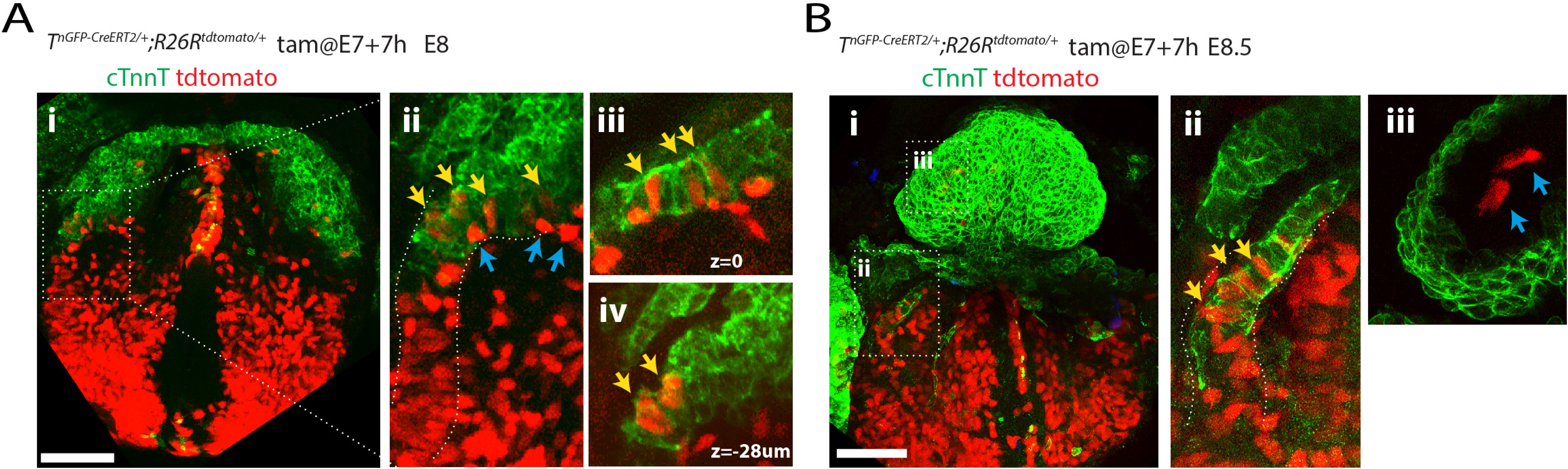
Descendants from late E7+7h T positive primitive streak cells contribute to posterior regions of the FHF. **(A-B)** Representative embryos resulting from the administration of tamoxifen at E7+7h in *T^nGPF-CreERT2/+^; R26R^tdTomato/+^* immunostained with cTnnT to reveal the cardiomyoyctes of the first heart field (green)(A) and heart tube (B). Yellow arrows identify tdTomato positive cardiomyocytes in the posterior first heart field (Aii-iv) and inflow tract (Bii). Blue arrows identify tdTomato positive endodermal cells (Aii) and endocardial cells (Biii). (Aii) shows magnified view of inset in (Ai). (Ai-ii and Bi-ii) are z-maximum projection, (Aiii-iv and biii) are single optical sections. Views are ventral. Single epicardial cells are labeled in each condition. Scale bar: 200 µm.

### Independent *Foxa2*-expressing cells located in the anterior primitive streak at the mid to late streak stages contribute to either the left or right ventricular myocardium

Genetic tracing experiments at E6-E6.5 (Frank et al., 2007; Bardot et al., 2017) showed that *Foxa2* positive cells contribute to cardiomyocytes in the ventricles but not in the atria. These results prompted us to test whether *Foxa2* positive cells contribute to ventricular myocardium at similar stages to the *T* expressing cells *i.e.* at the mid-late streak stage. We performed genetic tracing of the *Foxa2* positive cells using a *Foxa2^nEGP-CreERT2/+^* line expressing *CreERT2* downstream of the endogenous *Foxa2* in combination with the *R26R^tdTomato/+^* reporter line (Imuta et al., 2013) (Fig 3A-B). Mice were synchronized in estrus and timed matings performed for periods of 2 hours to better control embryonic stages.

**Fig 3.**
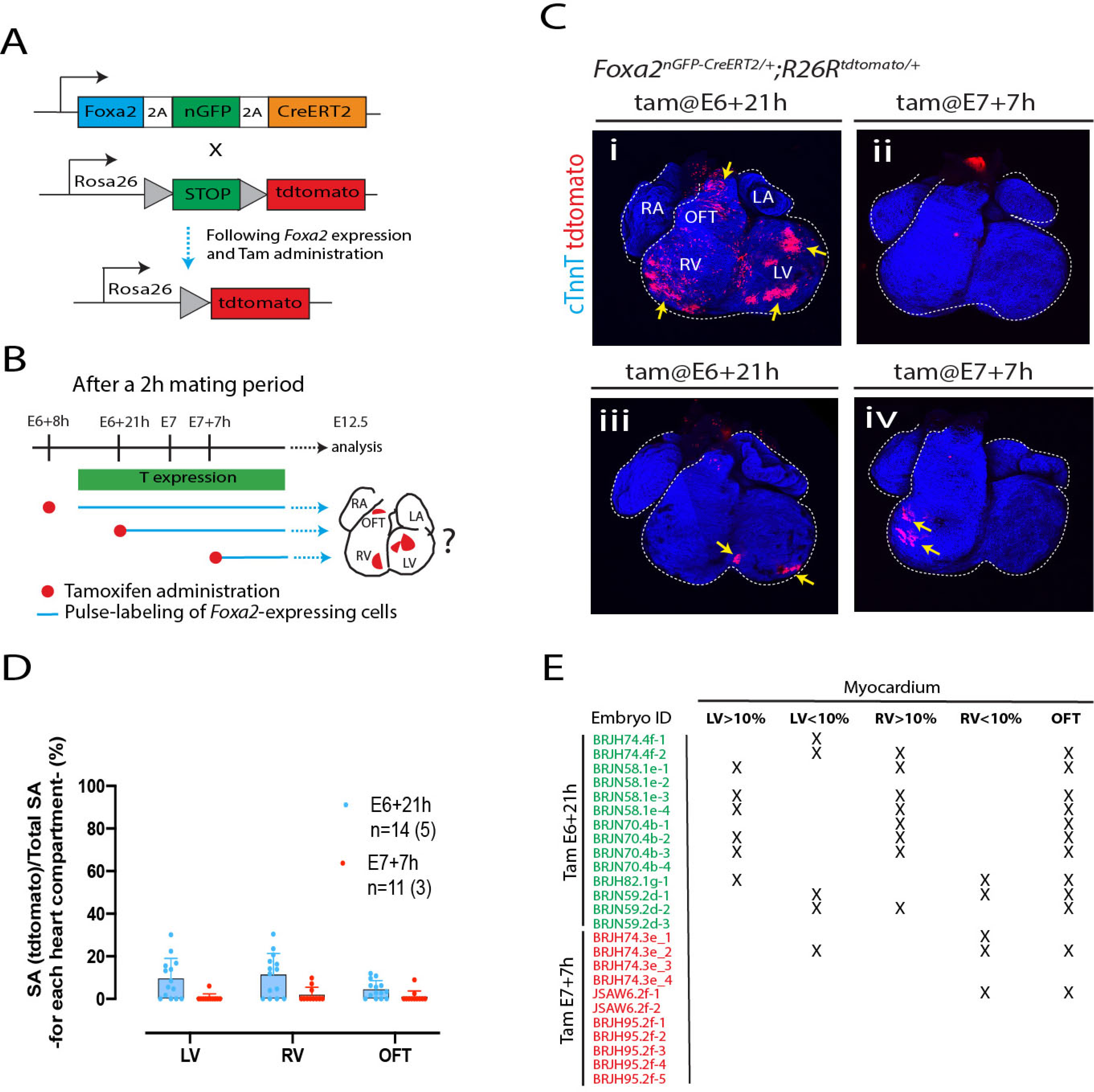
Independent *Foxa2+* progenitors contribute to the ventricles and outflow tract. **(A)** Schematics of the *Foxa2^nGPF-CreERT2/+^; R26R^tdTomato/+^* allele (Imuta et al., 2013). **(B)** Pulse-labelling of the *Foxa2*-expressing cells and their descendants, from stage (i) E6+21h, (ii) E7+7h by administration of tamoxifen to *Foxa2^nGPF-CreERT2/+^; R26R^tdTomato/+^* mice (0.08mg/body weight via oral gavage). Cell descendants in the heart analysed at E12.5. **(C)** Representative hearts resulting from the administration of tamoxifen at different time points in *Foxa2^nGPF-CreERT2/+^; R26R^tdTomato/+^* immunostained for cTnnT to reveal the cardiomyocytes (blue). Views are ventral. Single epicardial cells are also labelled in iii. **(D-E)** Summary of all hearts analysed. The contribution of the *Foxa2-*expressing cells to the different compartments of the heart is quantified by measuring the proportion of tdTomato positive myocardium. Numbers in brackets in (E) represent the number of litters assessed. Error bars are SD. Scale bar: 200 µm.

Administration of tamoxifen at E6+21h resulted in patches of tdTomato-expressing cardiomyocytes populating the left and right ventricle and outflow tract (Fig 3Ci). On average, the contribution of the tdTomato positive cells to the ventricles was low (left ventricle: 9.5% ± 9.5; right ventricle: 11.2% ± 10, p-value:ns). In 5 hearts out of 14, less than 10% of the left or right ventricle myocardium was populated by tdTomato-expressing cardiomyocytes. In addition, in 3 out of 14 heart no tdTomato-expressing cells could be detected in either the left or right ventricles (Figure 3D-E).

When we administered tamoxifen at E7+7h, tdTomato-positive cardiomyocytes could be detected in the left ventricle in only 1 out of 11 embryos and in the right ventricle in 3 out 11 embryos (Fig 3Cii, D and E). These cells covered less than 10% of the total left and right ventricle surface area (Fig 3D and E). In 3/14 hearts, small tdTomato-expressing domains formed either in the left (when labelled at E6+21h, n=2) or right ventricular myocardium (when labelled at E7+7h, n=1). This indicates that independent groups of *Foxa2* expressing progenitors exist in the primitive streak that contribute to the left and right myocardium (Fig 3Civ-v and E). We conclude that *Foxa2* positive primitive streak cells contribute to the ventricular myocardium at similar stages but in a lower proportion to the *T*-expressing cells.

Analysis of the embryos at stages when the cardiac crescent is differentiating into cardiomyocytes, confirmed that cells arising from *Foxa2* positive cells at E6+21h contribute to the myocardium in addition to other lineages including the epicardium and pericardium (S5A Fig.). This is consistent with previous studies showing that the FHF contribute to all these cell types (Bardot et al 2014, Devine et al 2014, Tyser et al 2021, Zhang et al. 2021). No contribution to the endocardium located below the presumptive myocardial epithelium was found in our conditions however (S5B Fig.).

These results prompted us to analyse the location of *Foxa2* and *T* expressing cells at the mid and late streak stages. Whole mount immunostaining for T and Foxa2 revealed coexpression in individual primitive streak cells at the mid-streak position of mid and late streak stages. Double positive cells are also detected in the definitive endoderm (Fig 4Bi-ii, Ci. and S6Ai-ii Fig) and at the distal tip of the embryo where the node forms (Fig 4Cii). These results indicate that *T* and *Foxa2* are not always in mutually exclusive populations of cells as previously reported (Burtscher and Lickert, 2009). Instead, it suggests *Foxa2* and *T* are co-expressed in a population of anterior primitive streak cells that contribute to the ventricular myocardium but not to the atria. Finally, Foxa2 is undetectable in the primitive streak from the OB stage onwards, the stages during which primitive streak cells switched their contribution from the ventricles to the outflow tract and atria (Fig 4E and F and S6D-D”i Fig).

**Fig 4.**
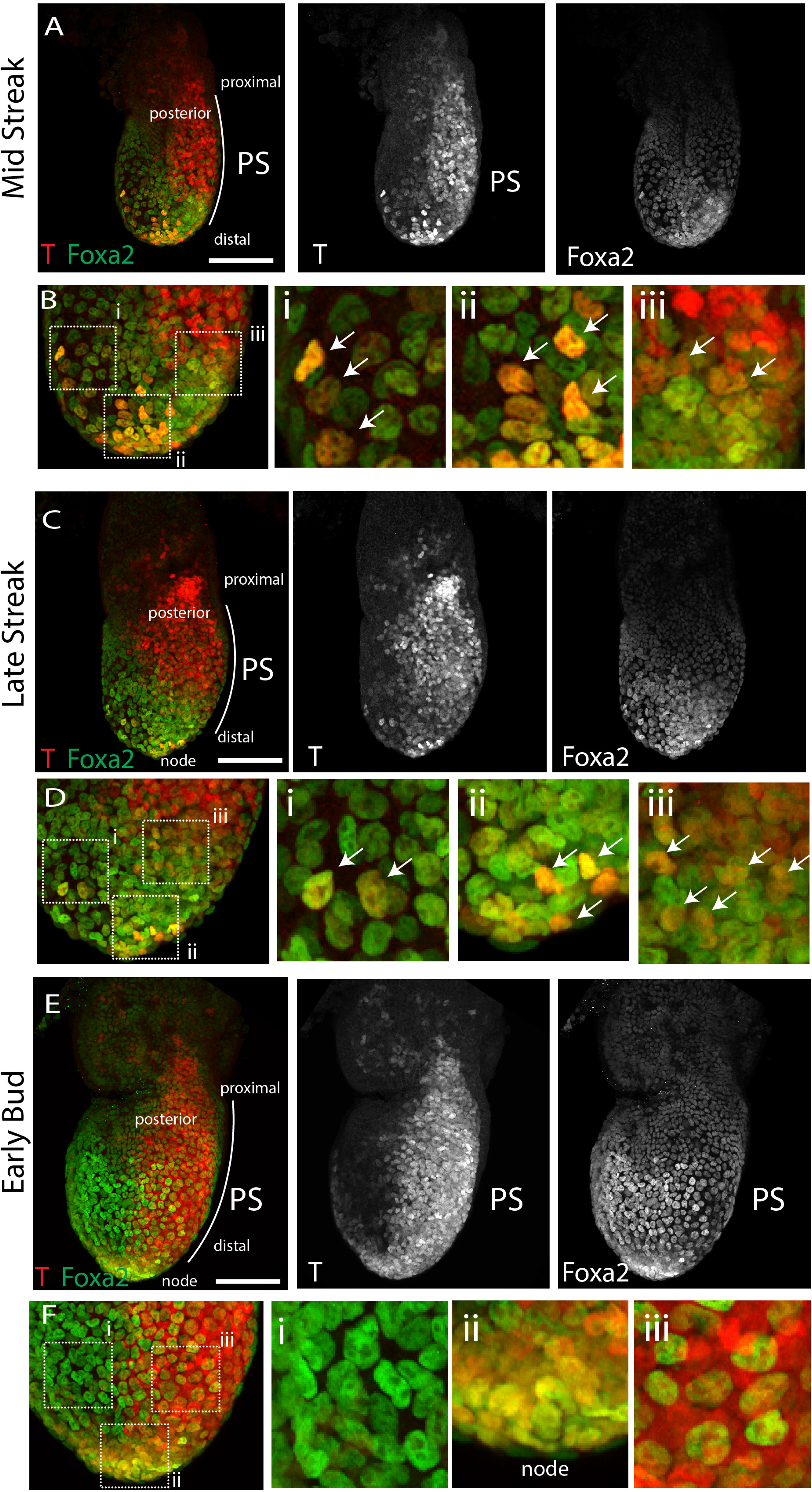
T and Foxa2 colocalise in primitive streak cells. Representative embryos of E6+21h MS (A-B) and LS (C-D) and E7+7h “early bud” EB (E-F). Embryos are immunostained for T (red) and Foxa2 (green). Images are Z-maximum projections. Views are lateral/slightly posterior to show the whole width of the primitive streak. Insets in Bi-iii, Di-iii and Fi-iii show magnified views and white arrows point to T+/Foxa2+ double positive cells. Scale bar: 100 µm.

### All ventricular progenitors derived from the *Foxa2* lineage express *Foxa2* at the mid-late streak stage

It is possible that a subset of the ventricular progenitors expresses *Foxa2* early and downregulate *Foxa2* and upregulate *T* by the mid to late streak stages. To test this idea, we administered tamoxifen in *Foxa2^nEGP-CreERT2/+^ R26R^tdTomato/+^* embryos earlier (E6+8h), collected the embryos at E6+21h and assayed for Foxa2 and T protein. Because of the variation in embryonic stages, we could analyse both the early and mid to late streak stages (Fig. 1F and G). As expected, at the early streak stage, we found Foxa2 positive cells with weak tdTomato signal located in the distal primitive streak (yellow arrows in Fig 5A). At the mid-streak stage, the streak had extended along its proximal-distal axis and we found tdTomato positive cells coexpressing Foxa2 and T in the anterior primitive streak (Fig 5B-G). We also found rare tdTomato positive cells expressing only T and not Foxa2 (red arrow in Fig 4Cii, yellow arrows in Fig 3F). Quantification of Foxa2 and T intensities revealed that the tdTomato positive cells located in the anterior primitive streak co-express low levels of Foxa2 and T (yellow arrows in Fig 5Ci-iii, Fig 5D-F and S7A Fig). The tdTomato positive cells located most distally expressed higher level of Foxa2 and low or no T (blue arrows in Fig 5Cii and iii and Fig 5D-F). These more distal regions correspond to where the axial mesoderm and definitive endoderm are established (Lawson et al 1991; Tam and Beddington, 1992). We conclude that cells expressing *Foxa2* early are not contributing to proximal regions of the primitive streak. Instead, these cells maintain *Foxa2* expression in the anterior primitive streak and contribute to the left and right ventricles at the mid-to-late streak stages (Fig 5G-H).

**Fig 5.**
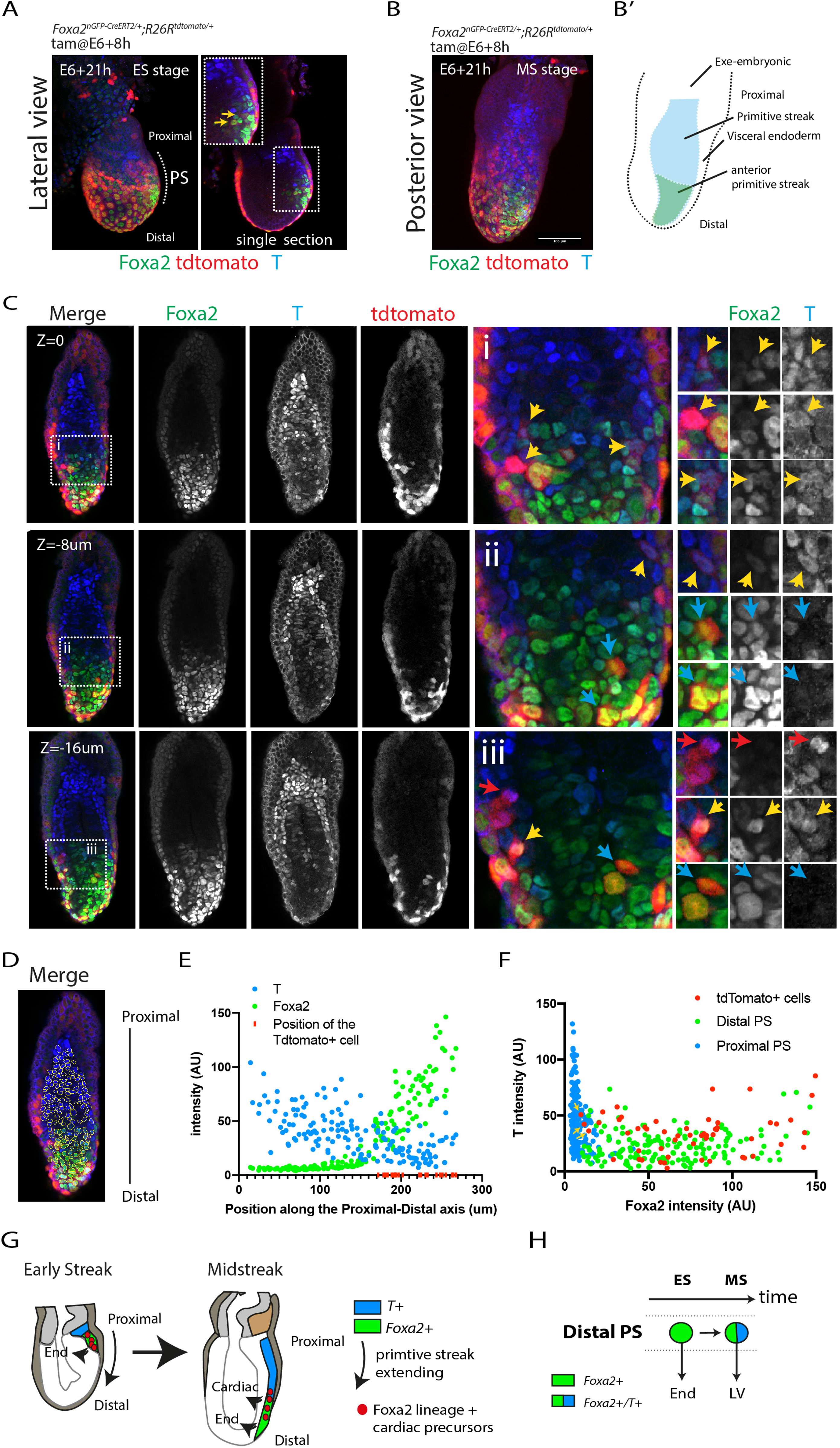
*Foxa2* lineage positive cells maintain *Foxa2* expression at the mid-late steak stage. **(A-B)** Representative embryos at the early (A) and mid-late (B) streak stages resulting from the administration of tamoxifen at E6+8h in *Foxa2^nGPF-CreERT2/+^; R26R^tdTomato/+^* immunostained for *Foxa2* (green) and T (blue). View is lateral in (A) and posterior in (B). (B”) Schematics of the embryo shown in (B). (**C**) Same embryo as in (B). Single optic sections are shown. (i-iii) are magnified views of the insets. Yellow arrows point to double Foxa2/T positive, blue arrows to Foxa2 positive T negative and red arrow to Foxa2 negative T positive tdTomato-expressing cells. (**D**) Representative example of a segmented image based on Foxa2 and T signal. (**E**) Quantification of T and Foxa2 signal intensities along the proximal-distal axis of the embryo in single cell nuclei. All the TdTomato positive cells are all located in the distal (anterior) portion of the primitive streak. (**F**) Quantification of Foxa2 and T intensities in single cell in distal (green), proximal (blue) primitive streak cells and tdTomato positive cells. (**G-H**) Model. As the primitive streak extend along the proximal-distal axis, from the early to mid-late streak stages, a subset of the *Foxa2* positive cells switch contribution from endoderm to cardiac (ventricles) progenitors and express *T*.

### Distal primitive streak cells downregulate *Foxa2* expression as they switch their contribution from the ventricles towards the outflow tract myocardium

Following tamoxifen administration in *Foxa2^nEGP-CreERT2/+^ R26R^tdTomato/+^* embryos at E6+21h, we found tdTomato-expressing cells located in the outflow tract myocardium (Fig 3Cii and Di and S5A Fig.). As previously mentioned, when we administered tamoxifen at E7+7h, no tdTomato-positive cardiomyocytes could be detected in the outflow tract (in 11 out of 13 hearts, Fig 3Ciii). This could indicate that a subset of the *Foxa2* positive primitive streak cells, at E6+21h, are outflow tract progenitors and not solely ventricular progenitors. Alternatively, primitive streak cells contributing to the outflow tract may arise from cells that had previously expressed *Foxa2* but no longer do so as the primitive streak switches its contribution from right ventricular to outflow tract myocardium by E7+7h. Consistent with this latter hypothesis, the presence of tdTomato-expressing cardiomyocytes in the outflow tract was always associated with the presence of tdTomato-expressing cardiomyocytes in the right ventricle (in 11/14 hearts, Fig 3E).

We next administered tamoxifen in *Foxa2^nEGP-CreERT2/+^ R26R^tdTomato/+^* embryos at E6+21h and fixed the embryos at E7+7h (Fig 6A). We immunostained embryos for Foxa2 and T protein. Computationally masking cells for Foxa2 expression (*i.e.* endodermal and axial mesoderm cells, Fig 6Aii-iii) revealed tdTomato-expressing mesodermal cells (*i.e.* cells that had expressed Foxa2) located in distal regions of the primitive streak and in the medial/dorsal mesoderm (Fig 6Aiii). No tdTomato positive cells were detected in lateral regions of the mesoderm.

**Fig 6.**
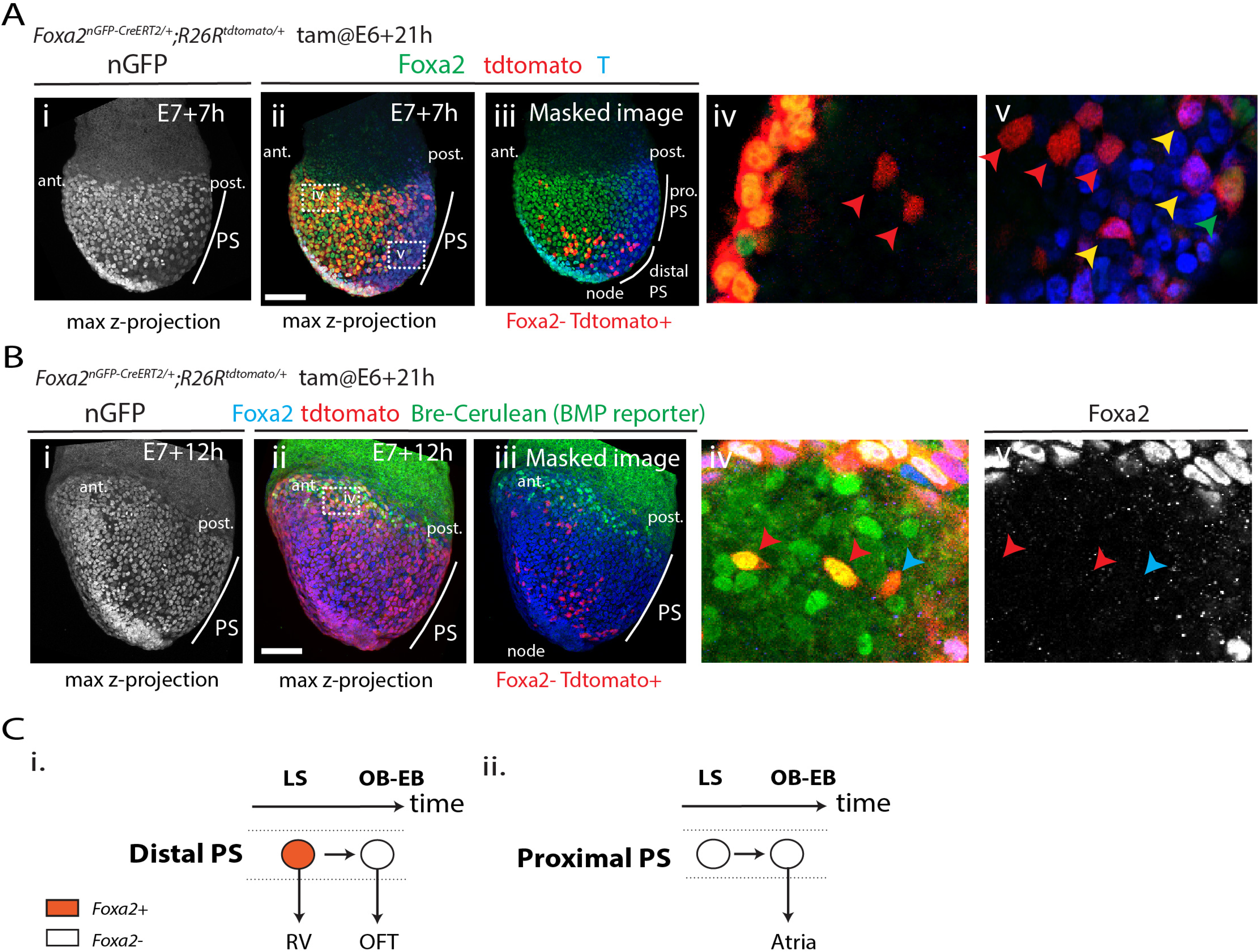
Ventricular precursors originate from distal regions of the primitive streak. **(A)** Localisation of the tdTomato+ cells from the *Foxa2* lineage are assessed (i-ii) at E7+7h -OB-EB stage-. nGFP signal from the transgene is shown in i. The embryos are immunostained for T (bleu) and Foxa2 (green) (ii). Image is a z-maximum projection. (iii) The same embryo (i-ii), but with Foxa2+ cells masked to reveal the tdTomato+/Foxa2-cells. Insets (iv) and (v) show magnified views in single optical sections of (ii). Red arrows point to tdTomato+/Foxa2-/T-mesodermal cells. Yellow arrows point to tdFtomato+/Foxa2-/T+ primitive streak cells. Green arrows point to tdTomato+/Foxa2+/T-endodermal cells. No tdTomato+/Foxa2+/T+ cells are identifiable. **(B)** Localisation of the tdTomato+ cells from the *Foxa2* lineage are assessed in Bre-cerulean (BMP reporter) embryos at (i-ii) E7+12h - EHF stage. nGFP signal from the transgene is shown in (i). The embryos are immunostained for Foxa2 (blue) (ii). Image is a z-maximum projection. (iii) The same embryo (i-ii), but with Foxa2+ cells masked to reveal the tdTomato+/Foxa2-cells. Insets (iv) and (v) show magnified views in single optical sections of (ii). Red arrows point to tdTomato+/Foxa2-/Bre-cerulean+ mesodermal cells. Blue arrow points to tdTomato+/Foxa2-/Bre-cerulean-cell. (**C**) (i) A common progenitor between the RV and the outflow tract in the distal primitive streak. *Foxa2* is downregulated as it switches its contribution toward outflow tract myocardium. (ii) Proximal PS contribute to the atria and are descended from cells that had not expressed *Foxa2* in their past. PS primitive streak, pro. PS: proximal primitive streak. Ant: anterior, Post: posterior. Scale bar: 100 µm

We then analysed all tdTomato-expressing cells within the distal primitive streak region and found these cells expressed T protein, (yellow arrows in Fig 6Av), although none co-expressed Foxa2 and T. Cells expressing Foxa2 but not T corresponded to endodermal cells, as indicated by their location in the outer layer of the embryo (green arrow in Fig 6Av). These experiments show that the outflow tract myocardium descendants from *Foxa2* progeny arise from the distal primitive streak. The outflow tract progenitors are *Foxa2* negative, but they arise from cells that expressed *Foxa2* previously. We propose that distal primitive streak cells downregulate *Foxa2* as they switch their contribution from the right ventricle to the outflow tract myocardium (Fig 6C). Atrial cells arise instead from the proximal primitive streak since they never arise from cells that had expressed *Foxa2* and all the cells located in the anterior primitive streak express *Foxa2* (Fig 5E and F).

We further characterised the tdTomato-expressing mesodermal cells arising from the *Foxa2* lineage. These cells have a cranial mesodermal identity since they express *Mesp1* (S8A Fig). In addition, they are located adjacent to endoderm expressing the nodal and Wnt antagonists Cerebus1 (Cer1) and Dkk1 respectively (S8B and D Fig.) (Nandkishore N. *et al*. 2018). The primitive streak and posterior mesoderm are instead marked by Wnt/β−catenin signalling activity (S8E Fig.). TdTomato positive cells located in anterior regions of the embryo (red arrows in S8C Fig) were unlikely to be endothelial cells since they did not express the endothelial marker Flk1 (for all the embryos we analysed (n=7). Instead, they are more likely to be cardiac progenitors since they were responding to BMP signalling (by E7+12h) and cardiomyocytes of the first heart field -or cardiac crescent- are characterised by high levels of BMP signalling activity at these stages (Klaus et al., 2007; Lescroart et al., 2018) (Fig. 6B and S9A-A” Fig.).

### Ventricular, outflow tract and Atrial progenitors have distinct locations and transcriptional profile within the heart fields

Having identified distinct population of *T*-expressing cells descendants, we reasoned their final locations could correlate with the previously described heart fields at gastrulation stages (Lescroart et al. 2018). We analysed the locations of the descendants of early E6+21h and late E7+7h *T*-expressing cells. These cells formed the ventricles and the majority of the outflow tract/atria, respectively (see above). We used a BMP reporter (S9A-A” Fig) and assayed for Phospho(P)-Smad1/5/8 to mark the first heart field and for Raldh2 to mark the posterior second heart field. Progenitors derived from the early E6+21h *T*-expressing cells contributed to anterior structures, distinct from the region occupied by late E7+7h *T*-expressing cells (Fig 7Ai-ii). The descendants from the late E7+7h were located in posterior mesoderm anterior to the node where Raldh2 localisation is strong (Fig 7Di-iv). They were instead mostly excluded from the first heart field domain in which BMP activity is high, and also from the cranial paraxial mesoderm with no detectable BMP activity (Fig 7Biii-iv, Ci-ii and S10A Fig). Some tdTomato-expressing cells were also sparsely intermingled within the posterior BMP positive domain and among these some had detectable BMP activity (see yellow and red arrows in Fig 7Bi-iv, Ci-ii and S10A Fig). This is consistent with a late recruitment of E7+7h progenitors to posterior regions of the cardiac crescent (see above).

**Fig 7.**
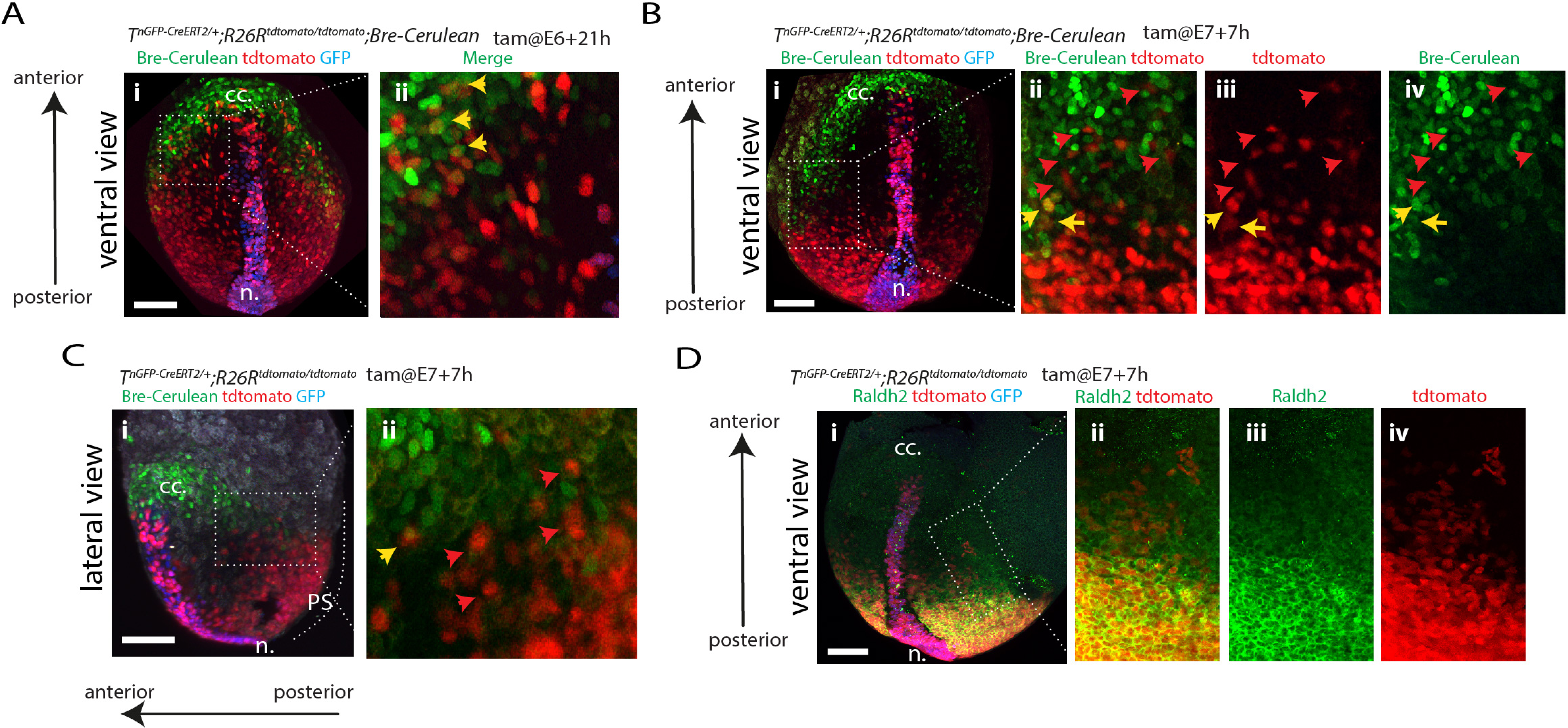
Ventricular, outflow tract and Atrial progenitors are located in distinct regions of the mesoderm. **(A-B)** TdTomato localisation in *T^nGPF-CreERT2/+^; R26R^tdTomato/tdTomato^, Bre-Cerulean* (BMP reporter) embryos at the EHF stage and following tamoxifen administration at E6+21h (A) and at E7+7h (B-C). Insets Aii, Bii-iv and Ci show magnified views of Ai, Bi and Ci respectively. Yellow arrows in Ai, Bii-iv and Cii point to double tdTomato/Bre-cerulean positive cells and red arrows in Bii-iv and Cii to tdTomato+ bre-cerulean-cells. (**D**) TdTomato localisation in *T^nGPF-CreERT2/+^; R26R^tdTomato/tdTomato^* embryos at the EHF stage, following tamoxifen administration at E7+7h and immunostained for Raldh2. Insets in ii-iv show magnified view of i. Images are z-maximum projection of 37 sections acquired every 5 µm and covering 185 µm. Interval between frames: 6mn and 30s. n: node, cc: cardiac crescent, ml: midline. EHF: early head fold. Scale bar: 100 µm

To ask whether molecular differences exist between the ventricular, outflow and atrial cardiac progenitors, we performed single-cell transcriptome analysis of four *T^nEGP-CreERT2/+^ R26R^tdTomato/tdTomato^* embryos dissected 7 hours after tamoxifen administration at E7+7h (early head fold -EHF-stage, Fig 8A). We reasoned we could use the presence or absence of tdTomato transcripts to discriminate the transcriptome of the tdTomato-positive outflow tract and atrial progenitors from the tdTomato-negative ventricular progenitors within the heart fields. We obtained 3494 high-quality single cell transcriptomes. We clustered cells into subpopulations, guided by available single cell transcriptomic data (S4A-C Fig) (Pijuan-Sala et al., 2019, E7+7h dataset, see Material and Method, S11A-D Fig and S12A-D Fig). Within these subpopulations, cardiac progenitors belonging to the previously defined first heart field -FHF-, anterior heart field -AHF- and posterior second heart field -pSHF- (Lescroart et al. 2018) and cranial paraxial mesoderm (Campione et al., 1999; Filipe et al., 2006; Nam et al., 2007; Zhao et al., 1996) were identified based on the expression of *Tbx5*, *Tbx20*, and *Hand2* for FHF; *Tbx1* and *Fgf8/10* for AHF; *Raldh2*, *Hoxb1and Tbx6* for pSHF; and *Pitx2*, *Alx1* and *Cyp26a1* for the cranial paraxial mesoderm (Fig 8C). Notably, *Fgf8* expression was reduced in the cranial paraxial mesoderm cluster, consistent with the absence of *Fgf8* expression in this cell population (Ilagan et al., 2006).

**Fig 8.**
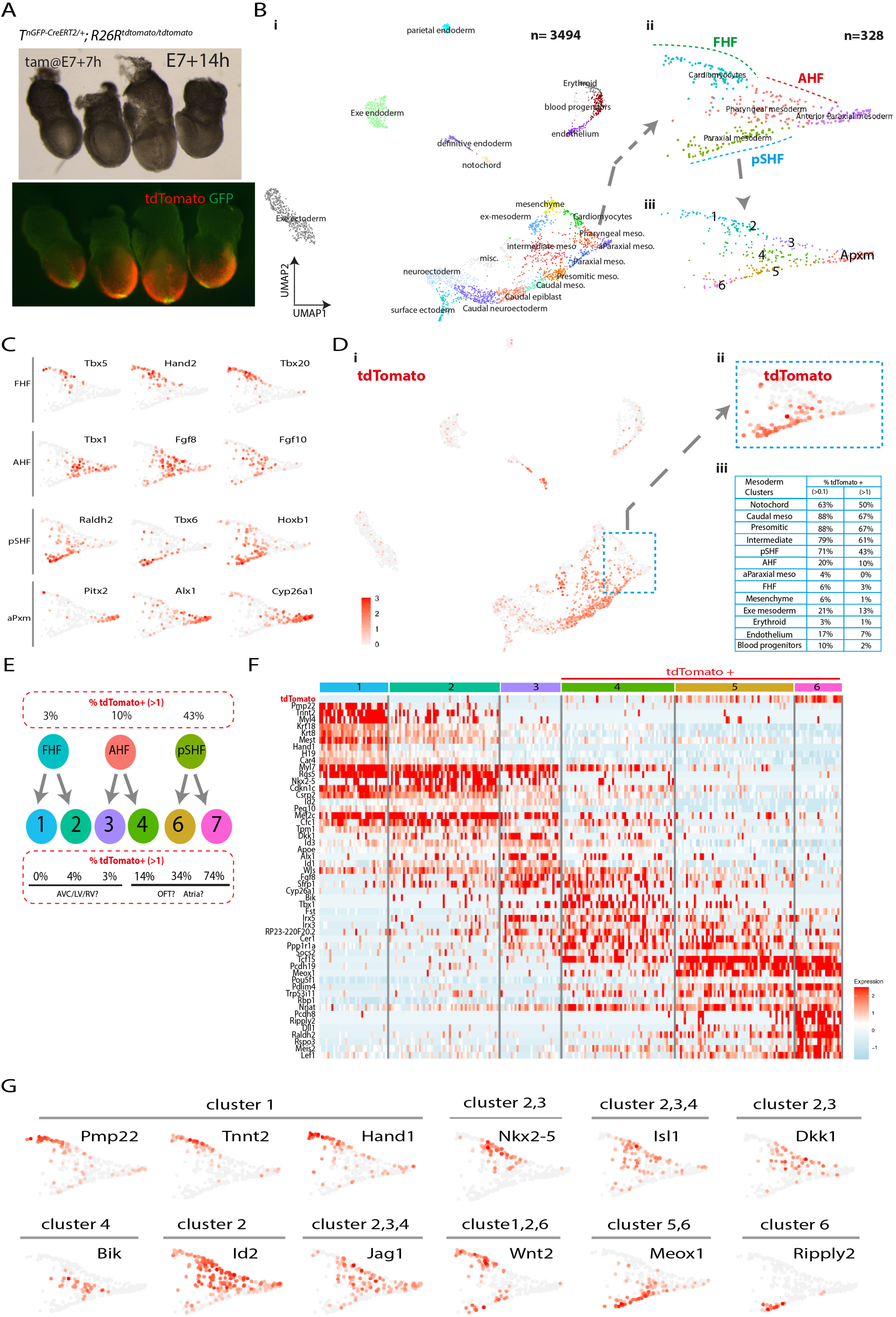
Subclusters corresponding to the left and right ventricle, outflow tract and atrial progenitors can be defined within the FHF, AHF and pSHF. **(A**) The four *T^nGPF-CreERT2/+^; R26R^tdTomato/tdTomato^* embryos (dissected at E7+14 h ∼EHF stage) analysed in the single cell transcriptomic assay. (**B**) UMAP plot showing the ectodermal, epiblast and mesodermal cell populations coloured by cluster identity (i). (ii-iii) Magnified view of (i) showing the cardiomyocytes, pharyngeal mesoderm, anterior paraxial and paraxial mesoderm clusters (ii) and subclusters (iii). The cardiomyocytes, pharyngeal mesoderm and paraxial mesoderm clusters are renamed FHF, AHF and pSHF based on the marker genes shown in (D). (**C**) UMAP showing the log normalised counts of the tdTomato gene (i). (ii) Magnified view of (i) showing the log normalised counts of the tdTomato gene in the FHF, AHF and pSHF clusters (**D**) UMAP showing the log normalised counts of the selected genes (**E**) Diagram showing the repartition of the tdTomato positive cells from the FHF, AHF and pSHF clusters into 6 subclusters, each with different proportion of tdTomato positive cells (see also S13A Fig). We hypothesis that cells in clusters enriched in tdTomato reads contribute to the OFT and Atria (**F**) Expression heat map of marker genes (source data Figure 5) and tdTomato. Scale indicates z-scored expression values. The full heat map with all the genes display is shown S13A Fig. (**G**) UMAP showing the log normalised counts of selected genes. n.: node, PS: Primitive streak, FHF: first heart field, AHF: anterior second heart field, pSHF: posterior second heart field, aPxm: anterior paraxial mesoderm, meso: mesoderm, Intermediate: Intermediate mesoderm, OFT: outflow tract, AHF: anterior heart field, pSHF: posterior second heart field, AVC: atrio-ventricular canal.

We next analysed the proportion of tdTomato positive cells in each cluster (S12C Fig, Fig 8Di-iii and S8_source data). Consistent with the imaging, we found that posterior mesoderm populations (*i.e.* caudal, presomitic, intermediate and paraxial mesoderm) and notochord were enriched in tdTomato positive cells (50-67%, scale data>1, Fig 8Ciii). Conversely, we found a lower frequency of tdTomato positive cells or none in anterior mesoderm populations including the anterior paraxial mesoderm (0%, scale data>1). Among the cardiac progenitors, the pSHF had the highest proportion of tdTomato-expressing cells (43%, scale data >1) followed by the AHF (10%, scale data>1). The FHF had very few tdTomato-positive cells (3%, scale data>1) (Fig 8Diii).

Since the AHF contribute to both the right ventricle and the outflow tract (Kelly et al. 2001), we next increased the resolution of the clustering to better distinguish tdTomato positive from tdTomato negative subpopulations. We observed the FHF, AHF and pSHF clusters could be subdivided into six discernible clusters (Fig 8E and F, S13A Fig and S8_source data). Three of these clusters were enriched for tdTomato-positive cells (14%, 34% and 74% for cluster 4, 5 and 6 respectively, scale data>1). The three other clusters were depleted of tdTomato-positive cells (0%, 4% and 3% for cluster 1, 2 and 3 respectively, scale data>1). We hypothesised that the tdTomato positive clusters included progenitors contributing to the outflow and atria. Conversely, we reasoned that the tdTomato negative clusters comprised progenitors derived from earlier *T*-expressing cells that contribute to the left and right ventricles and atrioventricular canal. We have summarised these results as a fate map for the different cardiac regions (Fig 11A-B) and we propose that left and right ventricular, outflow and atrial progenitors form molecularly distinct populations within the FHF, AHF and pSHF at the EHF stage mirroring their distinct origin in the primitive streak. We describe below the genes expressed in each of the tdTomato positive (4, 5 and 6) and negative (1, 2 and 3) subpopulations in relation to the known expression patterns in cardiac progenitors (Fig 8F-G and S7_source data res12 and Fig 8F).

Cells from cluster 5 and 6 -pSHF derived-expressed *Raldh2* and *Hoxb*1 (Fig 8C and F). Previous lineage tracing experiments have demonstrated that retinoic acid activated cells contribute to the outflow and atria (Dolle et al., 2010). Additionally, *Hoxb1* positive cells have been shown to contribute to the atria and to the inferior wall of the outflow tract that subsequently forms the myocardium at the base of the pulmonary trunk, in addition to patches of myocardium in the ventricles (Stefanovic et al 2020, Bertrand et al., 2011; Roux et al., 2015). Our data show *Tbx1* and *Raldh2* positive cells formed complementary cell populations within the tdTomato positive clusters. *Tbx1* is expressed in cluster 4 and 5 and Raldh2 in cluster 5 and 6. This is consistent with lineage tracing experiments demonstrating that *Tbx1* expressing cells within the AHF contribute to the outflow tract (Huynh et al., 2007). We also found that *Wnt2* was expressed in cluster 6. This is consistent with *Wnt2* expression in posterior mesoderm in addition to the anterior mesoderm where the prospective cardiac crescent forms (Costello et al., 2011; Monkley et al., 1996).

*Fgf8,* a known marker of the AHF (Lescroart et al. 2018), was enriched in both cluster 4 (low tdTomato positive) and cluster 3 (tdTomato negative). Lineage tracing experiments have shown that *Tbx1* marks part of the right ventricle progenitor domain within the AHF (Baldini et al., 2017; Huynh et al., 2007) while *Fgf8* marks it entirely (Crossley and Martin, 1995; Ilagan et al., 2006). Consistent with the hypothesis that cluster 3 corresponded to a right ventricular progenitor domain, we found *Tbx1* to be enriched within a subset of the cluster 3 (expressed in 30% of the cells) while *Fgf8* is expressed throughout (expressed in 70% of the cells). Also consistent with the hypothesis that cluster 2, 3 and 4 contained ventricular and outflow progenitors is the enrichment of *Jag1* in these three clusters, a gene shown to be expressed in progenitors derived from the *Foxa2* lineage (Bardot et al. 2020). *Jag1* is expressed anteriorly, complementary to the region where the cardiac crescent resides (Przemeck et al., 2003). *Dkk1*, a Wnt modulator controlling cardiac differentiation (David et al., 2008; Guo et al., 2019; Phillips et al., 2011), was also enriched in clusters 2 and 3. *Id 2,* which is essential for the specification of the heart tube-forming progenitors (Cunningham et al., 2017) is also enriched in clusters 2 and 3.

### The ventricular and outflow tract myocardium are prepatterned within the primitive streak

The signalling environment that cells encounter during migration influences the patterning of the heart fields (S13C Fig) (Andersen et al., 2018; Bardot et al. 2020; Guzzetta et al., 2020; Hochgreb et al., 2003; Klaus et al., 2007; Sirbu et al., 2008). However, our findings that the ventricles and the outflow/atria arise from distinct primitive streak cell populations raise the possibility that these progenitors are already molecularly distinct in the primitive streak. To address this question, we analysed the transcripts of cells from of tamoxifen treated *T^nEGP-CreERT2/+^ R26R^tdTomato/tdTomato^* embryos at the mid-late steak stages (3635 cells from nine embryos, Fig 6A) and OB-EB stages (3994 cells from four embryos, Fig 6A). Because we administered tamoxifen in these embryo, tdTomato positive mesodermal cells that had already emigrated could be visualised. In the mid-late streak stage embryos, tdTomato-positive cells were identified in the extra-embryonic mesoderm, consistent with the idea that the earliest progenitors to emerge from the primitive streak are contributing to the yolk sac blood cells and vasculature endothelium (Kinder et al., 1999). TdTomato-positive cells were also sparsely found in the embryonic mesoderm. In the OB-EB stage embryos, tdTomato-positive cells were found in the embryonic mesoderm where ventricular progenitors reside, and outflow and atrial progenitors are leaving the primitive streak at these stages. Unfortunately, the low level of expression at these times that meant very few tdTomato reads could be recovered and tdTomato positive cells could therefore not be identified in our single cell transcriptomic analysis at these early stages (15A, B Fig).

We clustered cells guided by available atlases of mouse gastrulation (Pijuan-Sala et al., 2019) (Fig 9B-C and S14A-C Fig). We analysed cells corresponding to the primitive streak and anterior primitive streak (aPS) clusters at the mid-late streak stages and cells in primitive streak cluster at the OB-EB stages (Fig 9E). The primitive streak cells at the mid-late streak and OB-EB stages did not express the *Cdx1/4* TFs associated with a posterior mesodermal identity. Nor did these cells express *Raldh2*, suggesting that the outflow and atrial progenitors start expressing *Raldh2* once the cells have reached their final location in the embryo (S14E Fig.).

**Fig 9.**
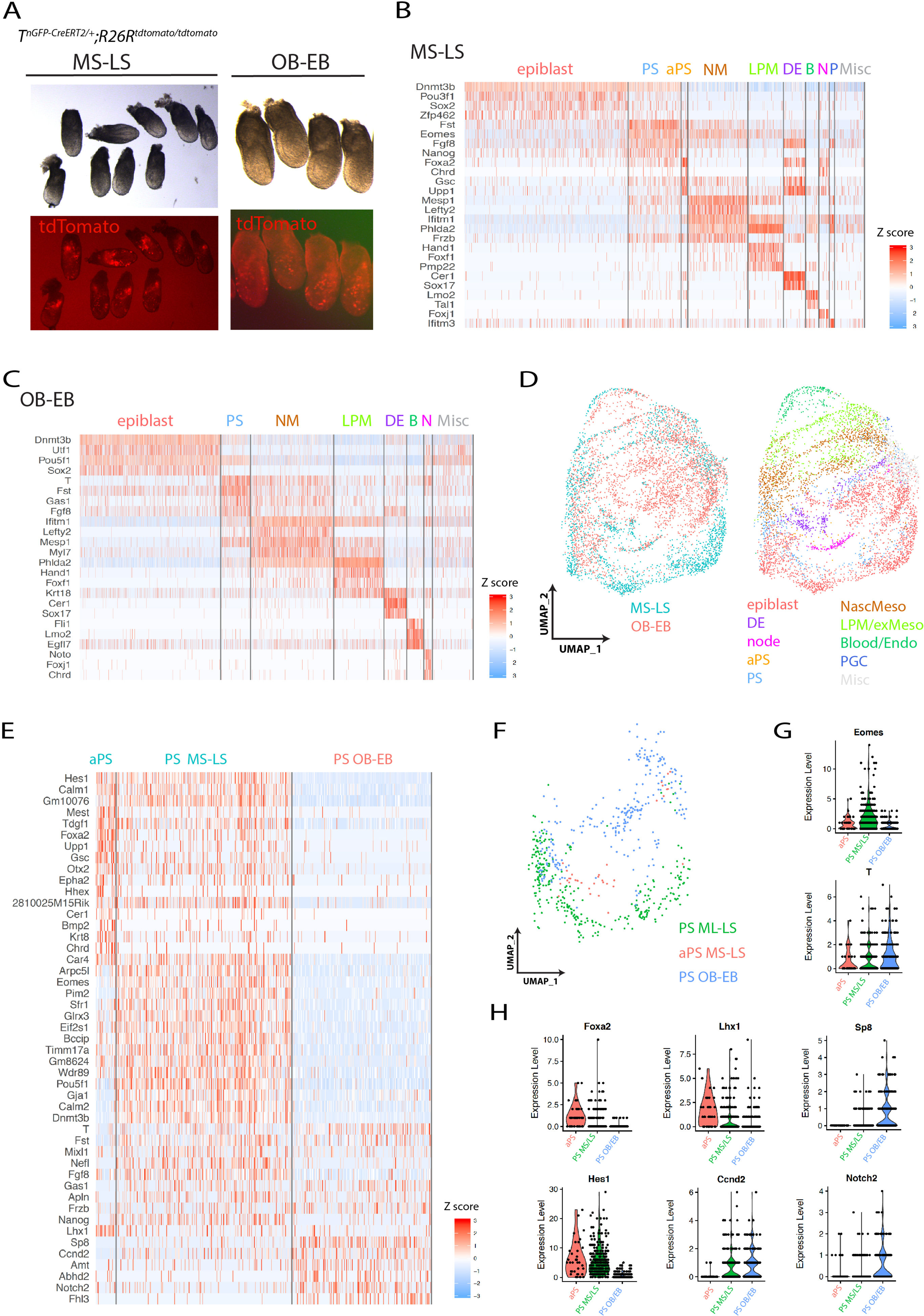
The molecular signature of the primitive streak cells contributing to the ventricles or outflow/atria are distinct. (**A**) Mid-late streak and OB/EB *T^nGPF-CreERT2/+^; R26R^tdTomato/tdTomato^* embryos analysed in the sc-RNA seq. Embryos were selected at the mid-late streak stages (dissected at E6+21h, and resulting from tamoxifen administration at E6+5h) and at the OB-EB stage (dissected at E7+3h, and resulting from tamoxifen administration at E6+21h). (**B-C**) Expression heat map of marker genes (source data Figure 6). Scale indicates z-scored expression values. (**D**) UMAP plot showing the integrated data from the two scRNA-seq mid-late streak and OB/EB datasets (see also Figure 6–figure supplement 1). Colour code correspond to the embryonic stage of collection or population identity. (**E**) Expression heat map of marker genes comparing the anterior primitive streak and primitive streak cells at the mid-late streak stages and primitive streak cells at the OB-EB stages (source data Figure 6). Scale indicates z-scored expression values. (**F**) UMAP plot of the anterior primitive streak, (aPS), mid-late streak (MS-LS) primitive streak and OB-EB primitive streak cells color coded. (**G-H**) Violin plots showing the normalized log2 expression value of *Eomes* and *T* (G) and *Foxa2*, *Lhx1*, *Sp8 Hes1*, *Ccnd2* and *Notch2* (H) in the anterior primitive streak (aPS) and primitive streak at mid-late streak and OB-EB stages. DE: definitive endoderm, aPS: anterior primitive streak, PS: primitive streak, Nascent meso: nascent mesoderm, LPM/Ex-meso: lateral plate mesoderm and Extra-embryonic mesoderm, mesenchyme. PGC: primordial germs cells. MS: mid-streak, LS: late streak, OB: no bud, EB: early bud.

Integration of the MS-LS and EB-OB datasets (Fig 9D) and differential gene expression analysis revealed major molecular differences between primitive streak cells contributing to either the ventricles or the outflow/atria (Fig 9E and F). The T-box transcription factor *Eomes* was strongly expressed in the mid-late streak stage primitive streak (ventricular progenitors) while its expression was lower at the OB-EB stages in the outflow/atrial progenitors (Fig 9G). This is in line with the function of *Eomes* in establishing the anterior mesoderm and specifying cardiac progenitors upstream of *Mesp1* (Costello et al., 2011). *T* had the opposite pattern of expression and showed a stronger expression at the OB-EB stages compared to the mid-late streak stages (Fig 9G). Fate mapping in the mouse also identified the anterior primitive streak adjacent to the organiser as a source of cardiac progenitors (Kinder et al., 2001) and *Hes1*, *hHex* (Thomas et al., 1998), *Gsc* (Blum et al., 1992), *Foxa2* (Ang and Rossant, 1994), *Upp1*, *Tdgf1* (Ding et al., 1998; Dono et al., 1993), *Bmp2, Epha2, Lhx1* (Costello et al., 2015; Shimono and Behringer, 1999)*, Otx2* (Acampora et al., 2003) and *Chrd* (Bachiller et al., 2000) were among the genes enriched in the anterior primitive streak at the mid-late streak stages (Fig 9E and H). *Sp8*, *Amt*, and *Notch2* (Williams et al., 1995) were preferentially enriched in the primitive streak at the OB-EB stages (Fig 6E and H). *Ccnd2* is expressed in the proximal regions of the primitive streak where the atrial progenitors are located (Wianny et al., 1998) and where BMP signalling activity is high (Morgani et al., 2018). *Mixl1* (Hart *et al*. 2002), *Fst* (Albano et al., 1994), *Gas1* (Lee and Fan, 2001), *Frzb* (Hoang et al., 1998) and *Apln* were expressed in the primitive streak at both the mid-late streak and OB-EB stages (Fig 9E). We conclude that ventricular, outflow and atrial progenitors derive from molecularly distinct groups of cells that occupy spatially and temporally discrete regions of the primitive streak.

### Live imaging reveals the migratory routes of the mesodermal cells

Finally, to confirm that early and late *T*-expressing cells migrated to distinct anterior-posterior locations within the heart fields, we tracked single cell in live *T^nEGP-CreERT2/+^ R26R^mg/+^* embryos (Fig 10E and S10_movie). Early mesodermal cells from distal locations (yellow arrows in Fig 10E) migrated anteriorly along the midline to regions where the first heart field is established. Conversely, both late distal and proximal mesodermal cells (orange and red arrows respectively in Fig 10E) contributed to more posterior regions of the mesoderm. The distal progenitors migrated to more medial regions (labelled as AHF, orange arrows in Fig 10E) and proximal cells to more lateral regions of the mesoderm (labelled as pCC and pSHF, red arrows in Fig 10E).

**Fig 10.**
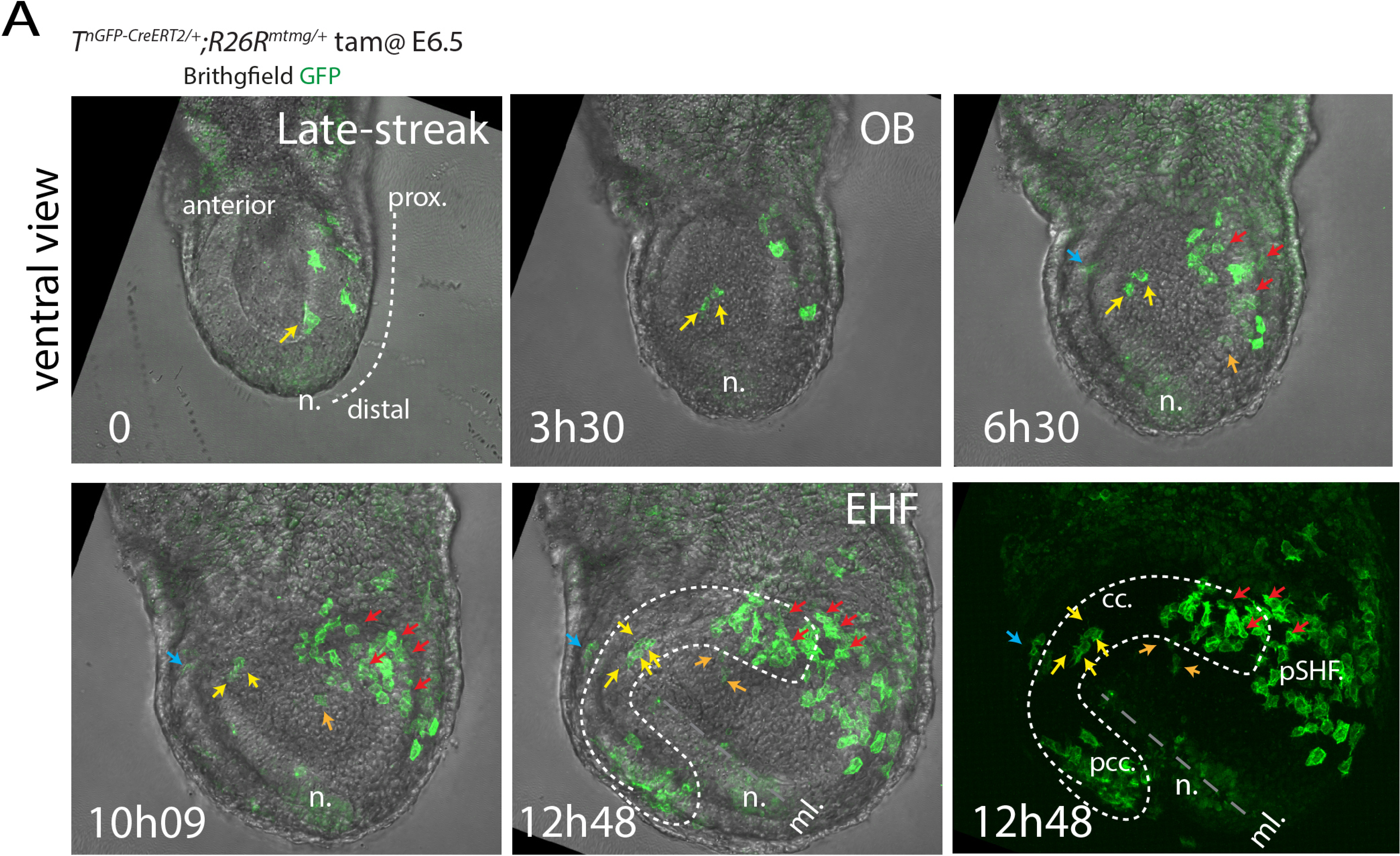
Live-imaging of the mesodermal cells reveal their trajectories during gastrulation. **(A)** Image sequence from time-lapse video (S9_movie) of an *T^nGPF-CreERT2/+^; R26^mgpf/+^* embryo resulting from the administration of tamoxifen at around E6.5 (overnight timed matings). Yellow arrows point to progenitor initially located in proximity to the node/ distal regions and migrating along the midline in medial regions of the prospective cardiac crescent. Red arrows point to progenitors initially located proximally and migrated towards posterior regions of the prospective cardiac crescent and posterior second heart field. Blue arrow points to a progenitor located at the embryonic-extraembryonic border. Images are z-maximum projection of 37 sections acquired every 5 µm and covering 185 µm. Interval between frames: 6mn and 30s. n: node, cc: cardiac crescent, pCC, posterior cardiac crescent, pSHF: posterior second heart field, ml: midline. EHF: early head fold.

## Discussion

Our findings, summarised in Fig 11A-C, reveal the temporal and spatial order in which different cardiac lineages arise within the primitive streak. The left ventricular progenitors are the first to leave the primitive streak at the mid-streak stage followed shortly after by the right ventricular progenitors at the late-streak stage. Progenitors contributing to the poles of the heart leave the primitive streak at the OB-EB stages. The outflow progenitors arise from distal regions of the primitive streak while the atrial progenitors are located in proximal regions of the primitive streak. These different subpopulations constitute molecularly distinct groups of progenitors within the heart field and the transcriptional differences in the primitive streak cells suggest that cardiac progenitors are prepatterned in the primitive streak prior to their migration. This organization of myocardial progenitors is conserved during evolution and in zebrafish, the ventricular and atrial myocardial progenitors are spatially separate in the blastula (Keegan et al. 2004).

**Fig 11.**
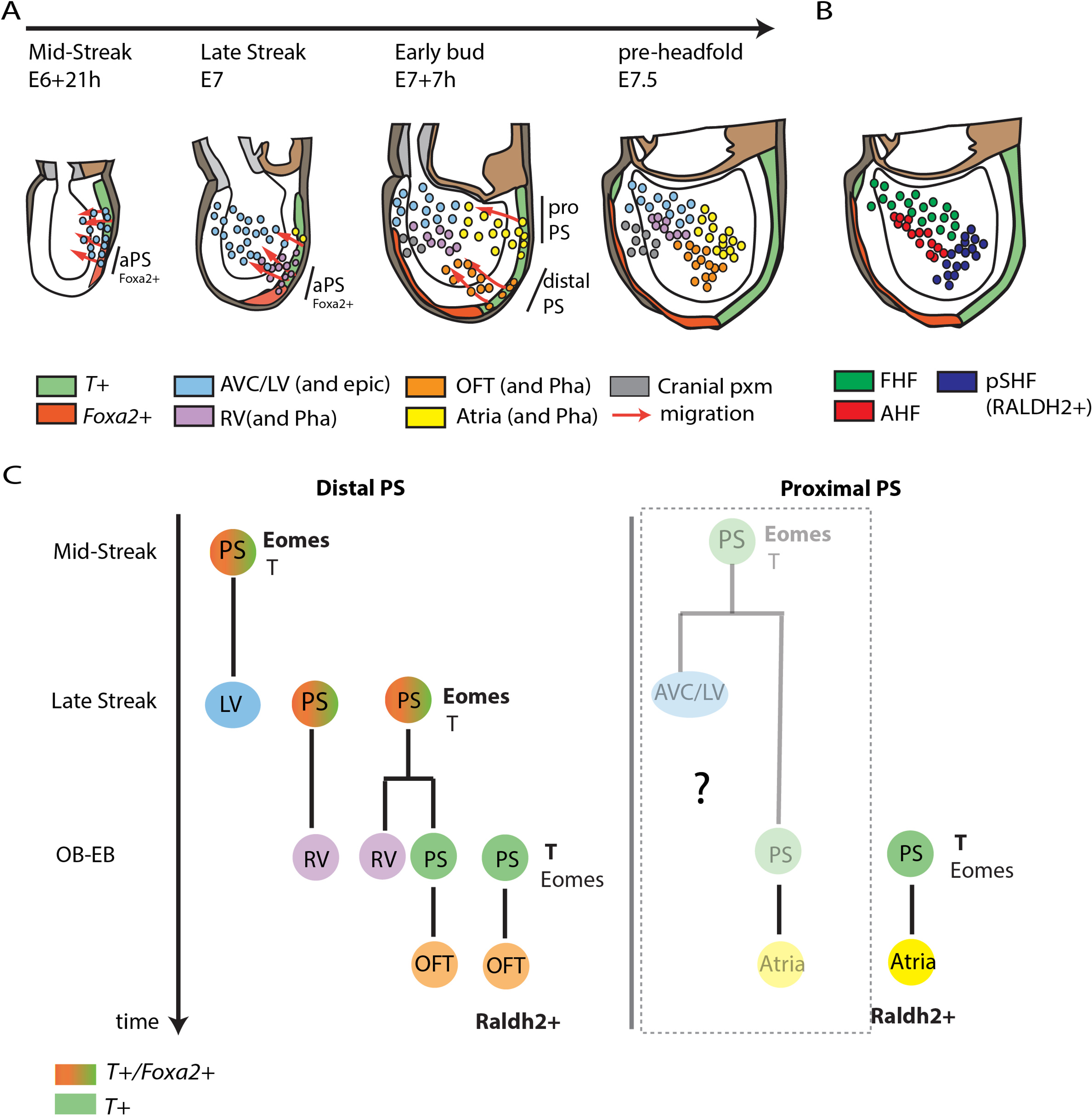
Model of early cardiac development. (**A**) Cells located in distal regions of the primitive streak contribute first to the left ventricle (mid-streak stage), then to the right ventricle (late-streak stage) and finally to the outflow tract (OB-EB stages). Although the outflow and atria leave the primitive streak at similar stages they arise from different regions. The outflow tract originates from distal locations in the primitive streak while atrial progenitors are positioned more proximally. The distal primitive streak cells express *Foxa2* when they contribute to the ventricles. They stop expressing *Foxa2* when they contribute to the outflow tract. (**B**) Proposed location of the FHF, AHF and pSHF in comparison to the hypothetical fate of these different cardiac regions at the pre-headfold stage in (A). (**C**) Lineage tree of the primitive streak. Within the distal end of the primitive streak, independent primitive streak cells contribute to the left ventricle, the right ventricle, the (at the mid-late streak stage) and to the outflow tract (OB-EB stages). A common progenitor between the right ventricle and the outflow tract exists. By contrast, atrial progenitors are located in the proximal primitive streak and a common progenitor between the left ventricle and atrium may exist in the primitive streak. LV: left ventricle, RV: right ventricle, OFT: outflow tract, Pha: Pharyngeal arches, epi: epicardium, cranial pxm: cranial paraxial mesoderm, FHF: first heart field, AHF: anterior heart field, pSHF: posterior second heart field. OB: no bud stage. EB: early bud stage, epic: epicardium.

### Primitive streak origin of the cardiac progenitors

While our *Foxa2* lineage tracing demonstrates an aPS origin of the ventricular myocardium, it is likely that more proximal regions of the PS marked by M*esp1* at the mid-late streak stage also contribute to the ventricles. Two lines of evidence support this hypothesis. First, the maximum contribution of the *Foxa2* progeny to the ventricles is 30-50% in *Foxa2* based lineage tracing experiments (Bardot et al. 2017 and this study). This suggests that the *Foxa2* progeny represent a subset of the whole pool of ventricular progenitors. Second, fate mapping analysis in the mouse demonstrated that some cells from the middle/proximal regions of the PS also contributed to the cardiac crescent or FHF (which mostly form the left ventricle) in addition to the aPS (Kinder et al 1998). Further investigation is required to characterize the descendants and contribution of the proximal primitive streak to the ventricles and whether or not they overlap with descendants of *Foxa2* progenitors. It is also unclear whether the *Foxa2* progeny contribute to a population of multipotent progenitors recently identified at the intra-/extra-embryonic boundary in the FHF (Zhang et al. 2021, Tyser et al. 2021). Instead, these progenitors may arise from an early population of proximal *Mesp1* positive primitive streak cells (Fig 11C) (Zhang et al. 2021).

Our *T*-lineage tracing experiments demonstrate that most of the atria arise from late OB-EB stages primitive streak. Our results further revealed that a subset of the atrial progenitors are gradually incorporated into posterior regions of the FHF to establish the inflow of the initial heart tube. This is consistent with the notion that the FHF contribute also contributes to part of the atria (Cai et al 2003, Später et al. 2014). The earliest PS cells contribute to the most anterior regions of the FHF which are fated to give rise mostly to the left ventricle and atrio-ventricular canal in addition to the epicardium in the heart while a rare contribution to the dorsal aspect of the right atria is noted (Zhang et al 2021). Thus, we propose that in the proximal regions of the primitive streak, progenitors contributing to the atrioventricular canal and left ventricle and are the first to emigrate followed rapidly by the atrial progenitors and a subset of these will contribute to posterior regions of the FHF (Fig 11C).

### Lineage relationships between cardiac progenitors

The *Foxa2* lineage tracing demonstrate the existence of a common lineage between the right ventricle and the outflow. This suggests a population of cells resides within the PS, the progeny of which contributes successively to the right ventricle and outflow tract progenitor pools while self-renewing (see model in Fig 11A-C). This is similar to a previous study that found the progeny of individually labelled primitive streak cells contributed to the notochord and somites while also leaving descendants in the primitive steak (Psychoyos and Stern, 1996). Live-imaging experiments will provide a more direct indication of the existence of asymmetric cell divisions mediating this process within the primitive streak.

While the right ventricle and outflow tract progenitors were identified in the anterior/distal primitive streak, atrial progenitors were located in the proximal primitive streak. This spatial segregation suggests that atrial progenitors are likely to constitute a pool of progenitors independent from right ventricle and outflow cardiac progenitors because of their physical separation. This is consistent with the original clonal analysis from Meilhac et al. (Meilhac et al., 2004), which concluded that atrial cells became clonally distinct before the presumptive right ventricle and outflow tract (Meilhac et al. 2004). Subsequent clonal experiments tracing *Mesp1* positive progenitors (Lescroart et al., 2014, Devine et al. 2014) also resulted in small clones spanning both the right ventricle and outflow tract compartments while independent clones were found in the atria (Lescroart et al., 2014, Devine et al. 2014) in addition to left ventricle-atrium clones (Devine et al. 2014). Thus, atrial progenitors are likely to constitute a pool of progenitors in the primitive streak distinct from a common progenitor for the right ventricle and outflow tract (see model in Fig 11C).

Meilhac et al. also observed two classes of larger clones extending across multiple compartments. This demonstrated the existence of two cardiac lineages in the embryo corresponding to the two heart fields (Meilhac and Buckingham, 2018; Meilhac et al., 2004). Our analysis and others (Devine et al. 2014, Lescroart et al. 2014 and Zhang et al. 2021) suggest that the segregation of two cardiac lineages could only occur before the onset of gastrulation, for example at the epiblast stage. A further subdivision into progenitors with more restricted cardiac lineages then arises by the time epiblast cells ingress through the primitive streak during gastrulation. This is reflected by the existence of progenitor subpopulations within the FHF and SHF corresponding to the prospective left and right ventricle, outflow tract and atria.

### A combination of intrinsic factors and inductive events specify and pattern the cardiac progenitors

Depending on the initial timing and site of ingression through the primitive streak, cardiac progenitors might adopt distinct migratory routes that exposes them to different signaling environments and influencing their cardiac fate. BMPs and Fgfs secreted by the anterior endoderm (Alsan and Schultheiss, 2002; de Soysa et al., 2019; Lough and Sugi, 2000; Schultheiss et al., 1997), retinoic acid expressed in the posterior lateral plate mesoderm (Hochgreb et al., 2003; Xavier-Neto et al., 1999) and signals expressed in the anterior intestinal portal (AIP) at later stages (Anderson et al., 2016) all contribute to the patterning of the cardiac progenitors. A Hedgehog-Fgf signaling axis has also been proposed to pattern the anterior mesoderm to allocate the head and heart during mesodermal migration (Guzzetta et al., 2020). Our analysis shows the primitive streak cells that contribute to either the ventricles or to the poles of the heart are molecularly distinct prior to migration. For example, changes in the transcriptional profile of primitive streak cells, including the downregulation of *Foxa2* expression, accompany the transition from contributing to the right ventricle to supplying the outflow tract. This mode of cellular diversification is reminiscent of temporal programmes in other tissues, such as neural progenitors (Doe, 2017), where sequential expression of distinct sets of transcription factors in progenitors defines the later differentiated cell types. These findings raise the possibility that cardiac progenitors are specified via a combination of both initial conditions set in the primitive streak and inductive events happening during migration (Bardot et al., 2020; Guzzetta et al., 2020) or once cells have reached their final location in the embryo.

Our analysis could not resolve putative differences between left and right ventricular progenitors in the primitive streak, and a more precise staging of embryos separating the mid from the late streak stages within a single cell transcriptomic assay will be required for this. Transcriptional differences between left and right ventricular progenitors may be established during migration by exposure to BMP and FGF signaling, both of which are known to mediate medio-lateral patterning of the mesoderm (Dorey and Amaya, 2010; Ilagan et al., 2006; Kelly et al., 2001; Tuazon and Mullins, 2015).

The observation that initial molecular differences in the primitive streak leads to the formation of groups of mesodermal cells that generate distinct populations of cardiac progenitors might inform *in vitro* cardiac differentiation protocols (Lee et al. 2017, Mendjan et al. 2014). We show that atrial cells are located in the proximal primitive streak where BMP signaling marked by P-Smad1/5/8 is high (Morgani et al., 2018). By contrast, ventricular progenitors originate in anterior primitive streak regions where cells are exposed to high Nodal signaling. This is consistent with methods to generate cardiac cells from human ESCs in vitro that rely on the exposure to different levels of Nodal and BMP signaling (Lee et al., 2017). A higher ratio of BMP4 to activin A signaling is required for the generation of Raldh2 positive mesoderm, which forms atrial cells. Conversely, the generation of ventricular cardiomyocytes relies on a higher level of activin to BMP4 signaling and the formation of Raldh2 negative CD235a positive mesoderm (Lee et al., 2017).

We conclude that cardiac progenitors are pre-patterned within the primitive streak and this prefigures their allocation to distinct anatomical structures of the heart. Further work will be required to test if generating the correct population of primitive streak cells can help to obtain purer populations of cardiomyocytes from pluripotent stem cells. Our analysis will also help to identify the initial mesodermal population that when dysregulated lead to specific malformations in the heart. For example, left ventricle hypoplasia results from a reduction in the number of specified cardiomyocytes within the mesoderm and this is due to a reduction in the expression of key cardiac transcription factors (Santos-Ledo et al., 2020). Whether such cardiac defects can arise from mutations affecting initial primitive streak cell populations and mesoderm remains to be addressed.

## Methods

### Experimental Model and Subject Details

All animal procedures were performed in accordance with the Animal (Scientific Procedures) Act 1986 under the UK Home Office project licenses PF59163DB and PIL IA66C8062.

### Mice

The *T^nEGFP-CreERT2/+^*(MGI:5490031) and *Foxa2^nEGP-CreERT2/+^* (MGI:5490029) lines (Imuta *et al*. 2013) were obtained from Hiroshi Sasaki. The *R26 ^Tomato Ai14^*^/ *Tomato Ai14*^ (Gt(ROSA)26Sor*^tm14(CAG-tdTomato)Hze^* (MGI:3809524), *Gt(ROSA)26Sor*^tm4(ACTB-tdTomato,-EGFP)Luo^, (MGI: 3716464) were obtained from the Jackson Laboratory. The *Mesp1^tm2(cre)Ysa^* (MGI:2176467) line was obtained from MRC Harwell, Mary Lyon. The TCF/Lef:H2B:mCherry WNT reporter line was generated in house by the Briscoe laboratory (Metzis et al. 2019).

### BRE:H2B-Turquoise BMP reporter line

The MLP:H2B-Turquoise sequence (originally from BRE-MLP-H2B-Turquoise pGL3 basic, Briscoe lab, unpublished) was cloned using In-Fusion Cloning (Takara, 639650) between two chicken insulators at the BamHI site of plasmid pJC5-4 (Chung et al., 1993) without the LCR fragment. The 1.6kb MLP:H2B-Turquoise fragment was amplified using Phusion High-Fidelity PCR Master Mix (ThermoFisher Scientific, F532L) according to the manufacturer’s instructions. Subsequently the 92bp BRE element was isolated with MluI/XhoI digest from BRE-MLP-H2B-Turquoise pGL3 basic and cloned 5’ to the MLP sequence of MluI/XhoI digested MLP:H2B-Turquoise pJC4-4 plasmid. The final plasmid was linearized with NdeI and used for pronuclear injection using fertilized embryos from the F1 hybrid strain (C57BL6/CBA). Mice with BRE reporter activity were verified by genotyping by a commercial vendor (Transnetyx). 5’-3’ Forward primer: CACAAGCTGGAGTACAACTACATCAGCGA Reporter 1: TCTATATCACCGCCGAC Reverse Primer: GGCGGATCTTGAAGTTGGCCTTGA. The BRE reporter line was maintained on the F1 background by crossing heterozygous *BRE:H2B-Turquoise* mice to F1 wild-type mice.

### Lineage tracing of the *T* and *Foxa2* expressing cells

Lineage tracing of *T and Foxa2* mesodermal progenitors was performed by crossing *T^nEGFP-CreERT2/+^* and *Foxa2^nEGP-CreERT2/+^* with *R26^Tomato Ai14/Tomato Ai14^* mice. To gain better control over embryonic staging, mice were synchronized in estrus by by introducing soiled bedding from a male’s cage into the females’ three days in advance. On the fourth day, mice were crossed over a two-hours period from 7am to 9am. Vaginal plugs were checked at 9 am and if positive, the embryonic day was defined at E0. This was followed by tamoxifen oral gavage with tamoxifen at 0.08mg/body weight (T5648 SIGMA) dissolved in corn oil at indicated times.

### Wholemount immunofluorescence and image acquisition

Embryos were fixed for 20min (for early E6.5-E7.5 embryos) or overnight (for late E12.5 hearts) in 2% PFA at 4°C, then permeabilized in PBST (PBS containing 0.5% Triton X-100) for 15min (for early E6.5-E7.5 embryos) or 1 hour (for late E12.5 hearts) and blocked for 5 hours (5% donkey serum, Abcam ab138579). Embryos were incubated overnight at 4°C with antibodies diluted in PBST (PBS 0.1% Triton X-100): rabbit anti-Estrogen Receptor alpha antibody (Sp1) (1:100, Abcam ab16660), rat anti-Flk1 (1:250, BD Biosciences 55307), goat anti-DKK1 (R&D Systems, AF1765), rabbit ranti-Aldh1a2 (Abcam ab96060), rat anti-CER1 (R&D Systems, MAB1986), goat anti-T (1:250, R&D Systems AF2085), rabbit anti-FOXA2 (1:250, Abcam ab108422); mouse anti-cTNT2 (1:250, Thermo Fischer Scientific Systems, MS295P0); rabbit anti-PhosphoSMAD1/5/8 (1:250, Cell Signalling D5B10 Rabbit mAb #13820). After washing in freshly prepared PBST at 4°C, embryos were incubated with secondary antibodies (Molecular Probes and Biotium) coupled to AlexaFluor 488 or 647 fluorophores and CF430 as required at 1:250 overnight at 4°C. Before imaging, embryos were washed in PBST at room temperature. Confocal images were obtained on an inverted Zeiss 710 confocal microscope with a 20X water objective (for early E6.5-E7.5 embryos) or a 10X air objective (0.4 NA) (for late E12.5 hearts) at a 1024 × 1024 pixels dimension with a z-step of 3-6-μm (2 × 2 tile scale, for the late E12.5 hearts). Embryos were systematically imaged throughout from top to bottom. Images were processed using Fiji software (Schindelin et al., 2012).

### Image Analysis

To segment the Foxa2 and T positive cells a Gaussian filter whose radius is adjusted to the typical size of a cell was first applied with Fiji (Schindelin et al., 2012). The resulting image was next converted to a mask by thresholding. When objects touched each other, a watershed on the binary mask. Finally, particle analyser generated a binary image with objects outside specified size (50) and circularity (0.75-1.00) removed. This process was repeated in each optical z-section of the z-stack. To quantify the surface area of the tdTomato labelled myocardial cells, patches of tdTomato cells were manually segmented and area measured with Fiji (Schindelin et al., 2012) for each individual heart within each litter.

### PCR analysis

Primers were designed to span the floxed SV40 poly(A) signal sequence removed from the genome following Cre recombinase-mediated recombination: 5’-3’ Forward primer: CGTGCTGGTTATTGTGCTGT; Reverse: CATGAACTCTTTGATGACCTCCTCGC. Primers yield a 1145 bp product from unrecombined DNA and a 274 bp product following recombination. gDNA was extracted using the HotSHOT method (Truett *et al*., 2000) from either the ear clip of an adult *R26 ^Tomato Ai14^*^/ *Tomato Ai14*^ mouse (unrecombined) or tail bud dissected from E9.5 embryos from a pregnant *T^nEGFP-CreERT2/+^ R26 ^Tomato Ai14^*^/ *Tomato Ai14*^ mouse, at 2, 4 and 12 hrs following oral gavage with Tamoxifen at 0.08mg/body weight (T5648 SIGMA) dissolved in corn oil. gDNA was then used in Q5 High-Fidelity 2X Master Mix (NEB) PCR reactions according to the manufacturer’s instructions. Amplicons were resolved on a 2% agarose gel with a 100 bp ladder.

### Sample preparation for single cell RNA sequencing

*T^nEGFP-CreERT2/+^ R26 ^Tomato Ai14^*^/ *Tomato Ai14*^ mouse embryos were imaged prior to dissociation with a Leica stereo fluorescent microscope with an exposure of ∼ 1mn to reveal the tdTomato signal. Sample preparation was done using previously established method (Delile et al. 2019). Mouse embryos were dissected in Hanks Balanced Solution without calcium and magnesium (HBSS, Life Technologies, 14185045) supplemented with 5% heat-inactivated foetal bovine serum (FBS). The samples were then incubated on FACSmax cell dissociation solution (Amsbio, T200100) with 10× Papain (30 U/mg, Sigma-Aldrich, 10108014001) for 11 min at 37°C to dissociate the cells. To generate a single cell suspension, samples were transferred to HBSS, with 5% FBS, rock inhibitor (10 μM, Stemcell Technologies, Y-27632) and 1× non-essential amino acids (Thermo Fisher Scientific, 11140035), disaggregated through pipetting, and filtered once through 0.35 μm filters and once through 0.20 μm strainers (Miltenyi Biotech, 130-101-812). Quality control was assayed by measuring live cells versus cell death, cell size and number of clumps and 10,000 cells per sample were loaded for sequencing.

### Analysis of scRNA-seq data

A suspension of 10,000 single cells was loaded onto the 10x Genomics Single Cell 3′ Chip, and cDNA synthesis and library construction were performed as per the manufacturer’s protocol for the Chromium Single Cell 3′ v2 protocol (10x Genomics; PN-120233) and sequenced on an Hiseq4000 (Illumina). 10x CellRanger (version 3.0.2) was used to generate single cell count data for each sample using a custom transcriptome built from the Ensembl mouse GRCm38 release 86 with the addition of the sequence from tdTomato Ai9 plasmid (Madisen et al., 2010). Due to 10x’s poly A bias, an additional 225 bases between the tdTomato gene stop codon and the bGH poly(A) signal was included to represent the WPRE gene. Depth of sequencing was 47975 mean reads per cell for the E7+14h dataset, 32629 mean reads per cell for the MS-LS dataset and 64340 mean read per cell for the OB-EB dataset. All subsequent analyses were performed with R (v.3.6.1) (R Core Team (2013)) using the Seurat (v3) package (Stuart et al., 2019). Primary filtering was performed on each dataset by removing from consideration: cells expressing unique feature counts fewer than 500, number of Unique Molecular Identifiers (UMIs) fewer than 1000 and cells for which mitochondrial genes made up greater than 3 times the standard deviation value of all expressed genes. Each dataset was normalised using the ‘LogNormalize’ function, with a scale factor of 10,000. The top 2000 highly variable genes were found using the ‘FindVariableGenes’ function and the data centred and scaled using the ‘ScaleData’ function. PCA decomposition was performed and after consideration of the eigenvalue ‘elbow-plots’, the first 50 components were used to construct Uniform Manifold Approximation Projection (UMAP) plots. Samples E7+14h and E7.75 from Pijuan et al. (Pijuan et al 2019) were integrated using Seurat standard integration workflow. Common anchor features between both experiments were selected using the ‘FindIntegrationAnchors’ functions, using the first 20 dimensions. Cluster identity from Pijuan et al. E7.75 dataset was next transferred to the E7+14h dataset. Cluster specific gene markers were identified using the ‘FindMarkers’ function using the settings (min.pct=0.25; min.diff.pct = 0.1 and return.thresh=0.0001), which uses the Wilcoxon rank sum test to compare each cell belonging on one cluster versus all other cells. Genes were ranked based on logFC and the highest logFC was used to determine unique markers per cluster. Selected cluster marker genes were used to draw a heatmap showing the expression of tdTomato on the top row (S12A Fig and S7_source data _res4.5). The clustering analysis was then repeated by increasing the resolution parameter and discernible clusters with marker genes corresponding to cluster 1-6 and anterior paraxial mesoderm were identified (S7_source data_res12). Cluster specific markers were also identified, after subsetting only clusters, 1, 2, 3, 4, 5 and 6 (Fig 8F and S7_source data_Fig 8F). Samples E6+21h and E7+3h were similarly integrated using Seurat standard integration workflow. Common anchors feature between both experiments were selected using the ‘FindIntegrationAnchors’ functions, using the first 20 dimensions. Clusters were labelled and grouped using pre-existing cell markers (Pijuan et al 2019) and differential expression between clusters were determined using the “FindMarkers Function”, without logfc or minimum percentage expressed thresholds. Genes showing the largest logFC and smallest adjusted p value between clusters were used to generate heatmaps (S9_source data).

### Embryo culture and two-photon live-imaging

Embryos were dissected at E6.5 in pre-equilibrated DMEM supplemented with 10% foetal bovine serum, 25 mM HEPES-NaOH (pH 7.2), penicillin (50μml21) and streptomycin (50mgml21). Embryos were cultured in 100% fresh rat serum filter sterilized through a 0.2 mm filter. To hold embryos in position during time-lapse acquisition, we made bespoke plastic holders with holes of different diameters (0.3–05 mm) to ensure a good fit of embryos similarly to the traps developed by Ivanovitch et al. (2017) and Nonaka et al. (1999).

Embryos were mounted with their anterior side facing up. To avoid evaporation, the medium was covered with mineral oil (Sigma-Aldrich; M8410). Before starting the time-lapse acquisition, embryos were pre-cultured for at least 2 hours in the microscopy culture set up. The morphology of the embryo was then carefully monitored and if the embryos appeared unhealthy or rotated and or moved, they were discarded, otherwise, time-lapse acquisition was performed. For the acquisition, we used the multi-photon MPSP5 equipped with a 5% CO2 incubator and a heating chamber maintaining 37°C. The objective lens used was a HCX APO L 20x/1.00 W dipping objective, which allowed a 2mm working distance for imaging mouse embryos. A SpectraPhysics MaiTai DeepSee pulsed laser was set at 880 nm and used for one-channel two-photon imaging. Leica Las AF software was used for aquisition. Image settings was: output power: 250 mW, pixel dwell time: 7μs, line averaging: two and image dimension: 610 × 610 μm (1024 × 1024 pixels). To maximize the chance of covering the entire embryo during the long-term time lapse video, we allowed 150–200 μm of free space between the objective and the embryo at the beginning of the recording.

### Data availability

Single cell RNA sequencing data have been deposited in NCBI under the accession number GSE153789.

## Supporting information

S1_raw image

S2_raw image

## Acknowledgements

The authors would like to thank the Science Technology Platforms at the Francis Crick Institute. In particular, we thank the Advanced Light Microscopy facility, the Advanced Sequencing Facility, Bioinformatics and Biostatistics Facility and the Biological Research Facility for their ongoing support and access to equipment. We are grateful to Robert Goldstone and Amelia Edwards for excellent support with single cell sequencing, Teresa Rayon for assistance with single cell preparation, Gavin Kelly for support with single cell data processing and Joe Brock for research illustration. We thank Teresa Rayon, Florencia Cavodeassi and Peter Scambler for comments on the manuscript and members of the Smith and Briscoe lab for useful discussion. K.I. has received funding from HFSP LTF (LT000609/2015-L). Work in the J.C.S. and J.B.’s lab was supported by the Francis Crick Institute, which receives its core funding from Cancer Research UK (FC001-157, FC001-051) the UK Medical Research Council (FC001-157, FC001-051), and the Welcome Trust (FC001-157, FC001-051). Work in the J.B.’s lab was also supported by the the European Research Council under European Union (EU) Horizon 2020 research and innovation program grant 742138.

## Author contributions

K.I.: Conceptualization, Investigation, Formal analysis, Visualisation, Writing – Original Draft Preparation. P.S.B. and P.C.: Formal analysis, J.D.: Data curation, N.M.G., D.S.: Investigation, R.A.J.: Investigation, Visualisation, D.M.B: Methodology, J.C.S.: Funding acquisition, Supervision, Writing-Review & Editing, J.B.: Funding acquisition, Conceptualization, Supervision, Writing-Review & Editing.

## Supporting information

**S1 Fig.**
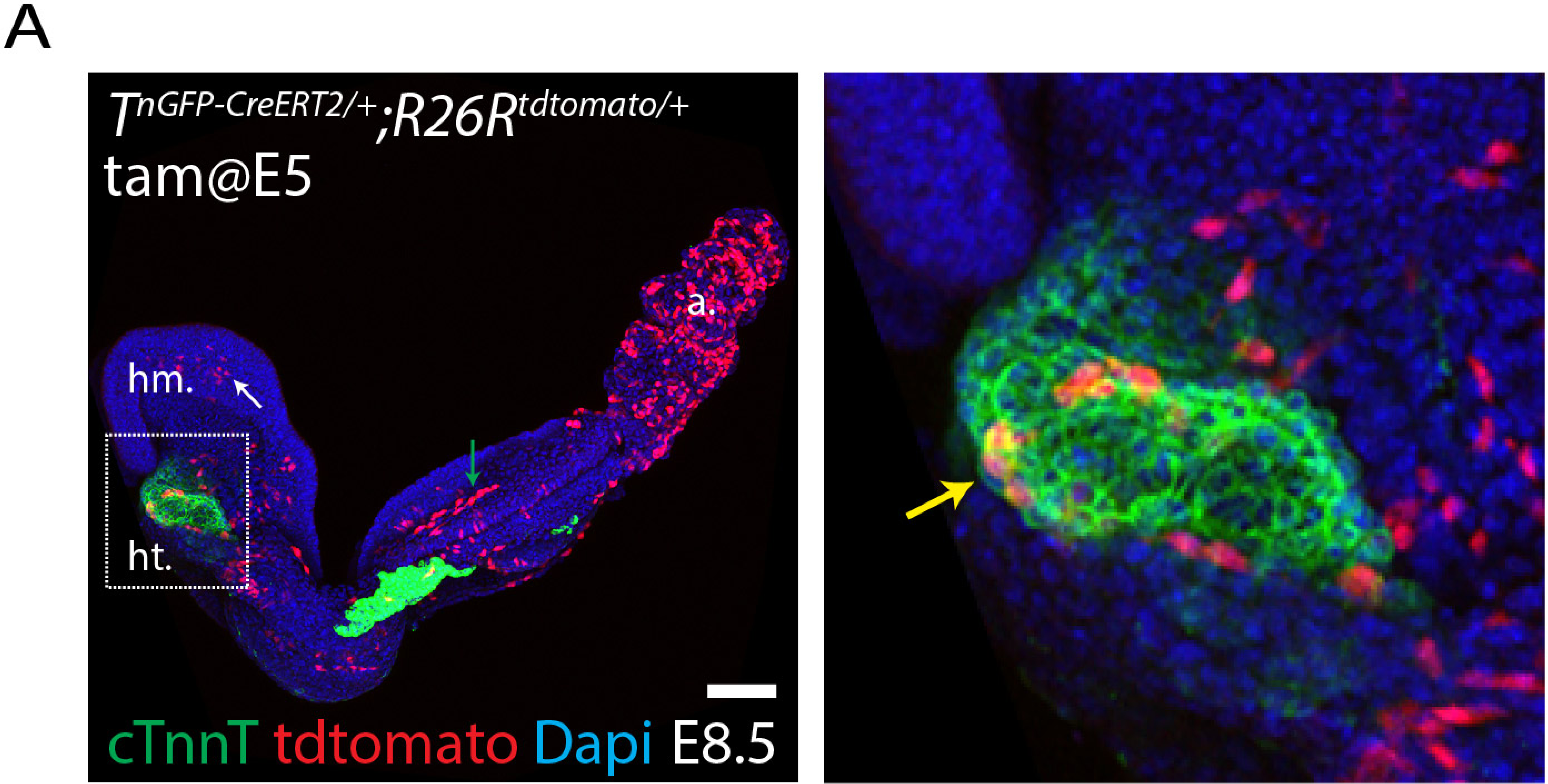
Tamoxifen activity persists for at least 24 hours when administrated at a high dose by oral gavage. **(A)** The administration of a high dose of tamoxifen (0.08mg/bw by oral gavage) at E5 in *T^nGPF-CreERT2/+^; R26R^tdTomato/+^* mice leads to the presence of tdTomato positive-cells in mesoderm derivatives including cardiomyocytes (see yellow arrow in inset), head mesenchyme (red arrow), endothelium (green arrow) and allantois. hm: head mesoderm, ht: heart tube, a: allantois. Mouse were mated for a two-hour period. Scale bar: 100 µm

**S2 Fig.**
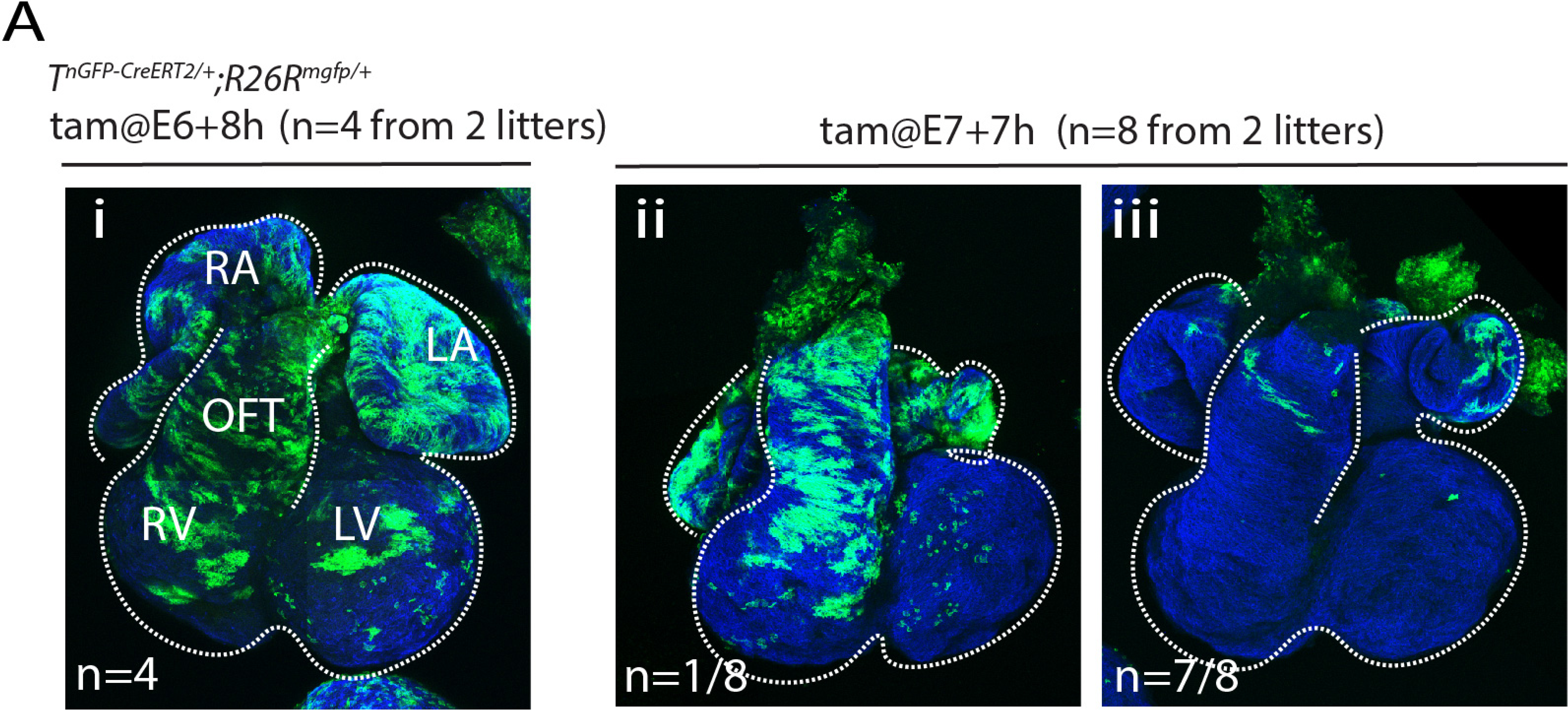
Genetic tracing of the *T+* primitive steak cell with the R26mt/mg reporter. **(A)** Representative hearts resulting from the administration of tamoxifen at E6+8h (i) and E7+7h (ii-iii) in *T^nGPF-CreERT2/+^; R26R^mtmg/+^* immunostained with cTnnT to reveal the cardiomyoyctes (blue).

**S3 Fig.**
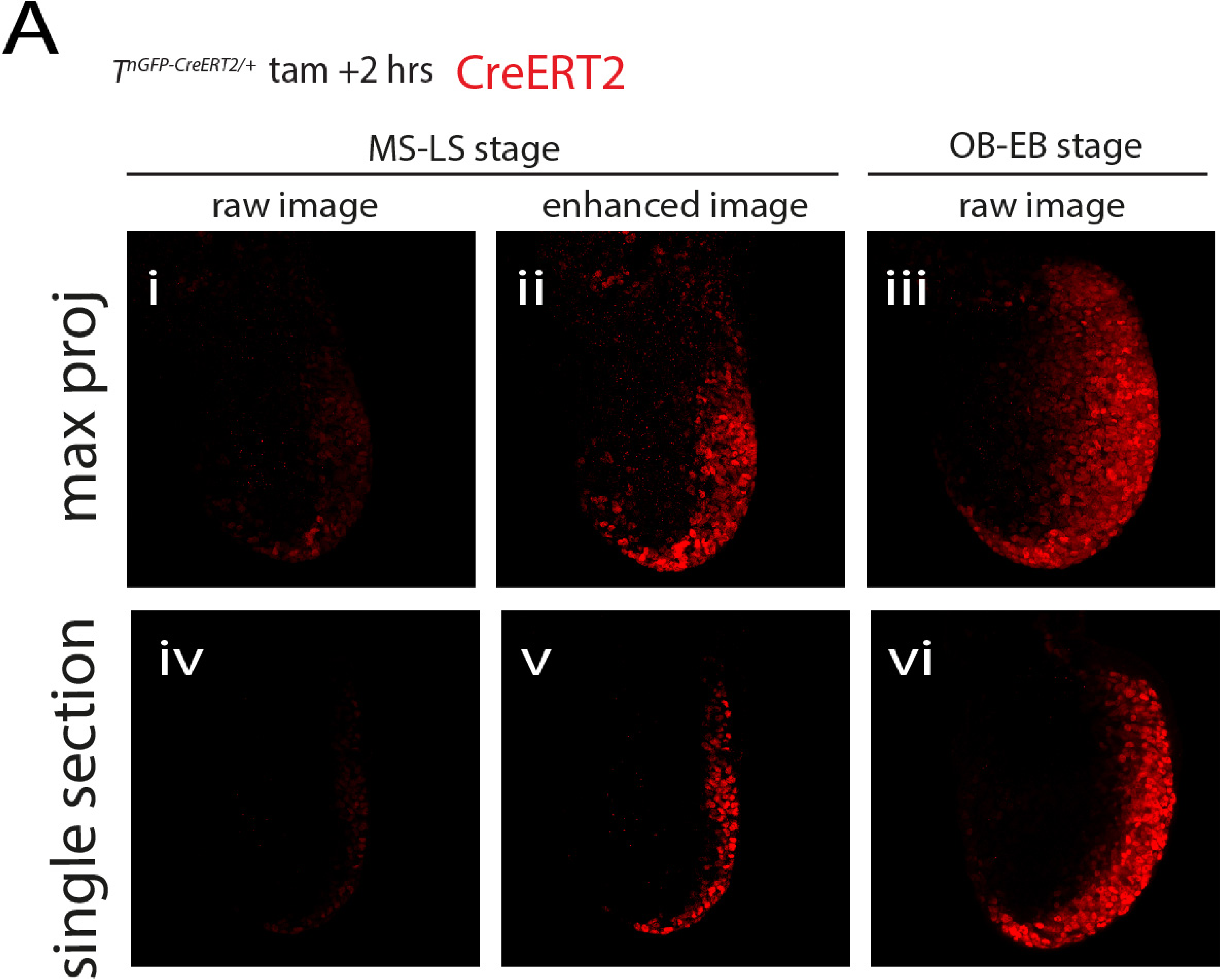
CreErt2 nuclear localisation 2 hours after tamoxifen administration. **(A)** Representative embryos resulting from a 2-hours pulse of tamoxifen via oral gavage (0.08mg/bw) immunostained with eostrogen receptor. Embryos have been immunostained simultaneously and image under the same conditions. Maximum z-projection (i-iii) and single optical sections (iv-vi) are shown.

**S4 Fig.**
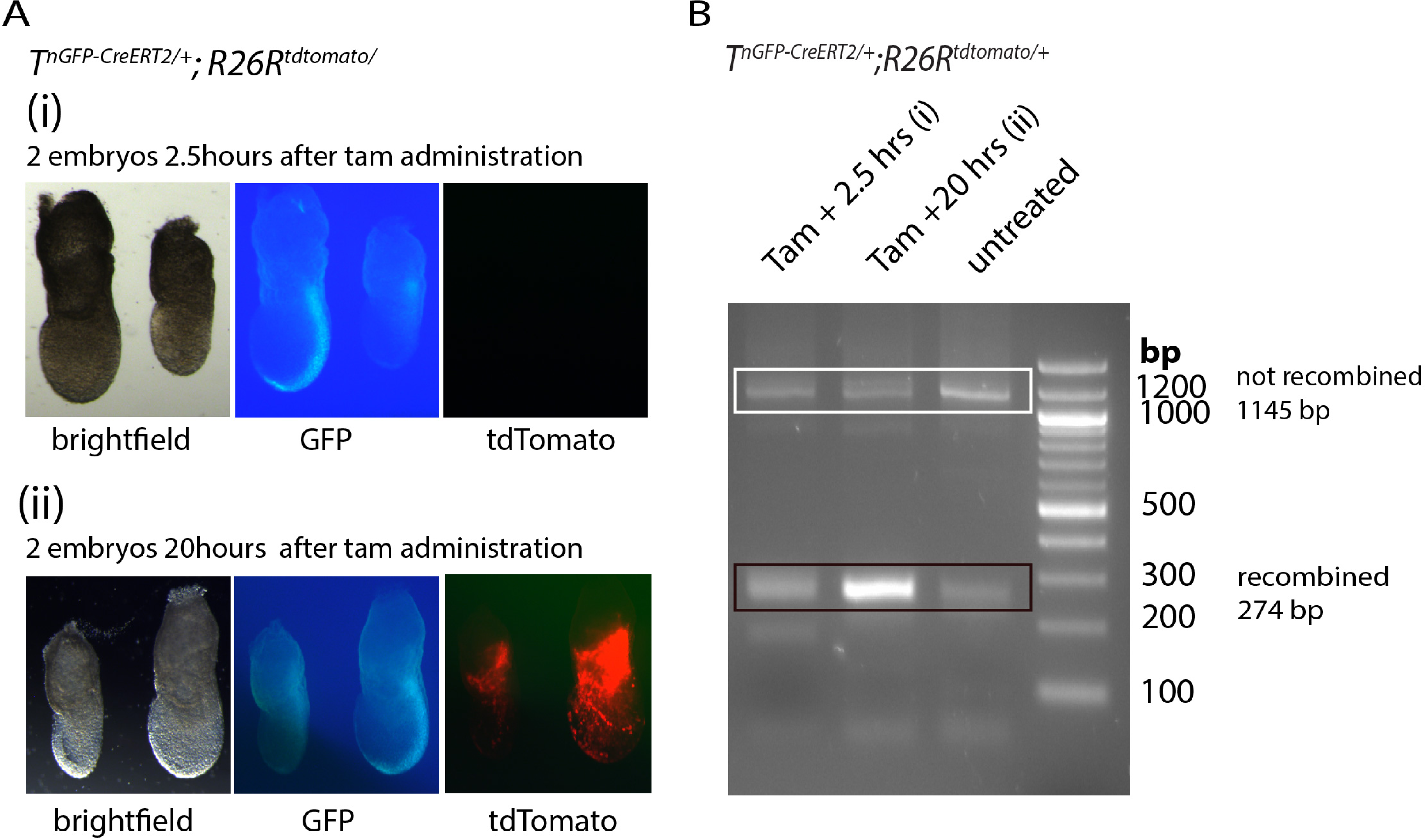
Recombination of the R26RtdTomato reporter is occurring 2.5 hours after tamoxifen administration by oral gavage. **(A-B)** PCR amplicons generated from the genomic region in which Cre-mediated recombination occurs from *T^nGPF-CreERT2/+^; R26R^tdTomato/tdTomato^* s embryos (A), resolved on an agarose gel (B). Before recombination, the PCR product is 1145 bp (white rectangle); after recombination it is 274 bp (black rectangle). Template gDNA was extracted from either an ear clip of an adult *T^nGPF-CreERT2/+^; R26R^tdTomato/tdTomato^* mouse (untreated) or *T^nGPF-CreERT2/+^*; *R26R^tdTomato/tdTomato^* embryos (i-ii) following oral gavage with Tamoxifen, as labelled. An increase in the proportion of the recombined band can be seen over time following Tamoxifen administration.

**S5 Fig.**
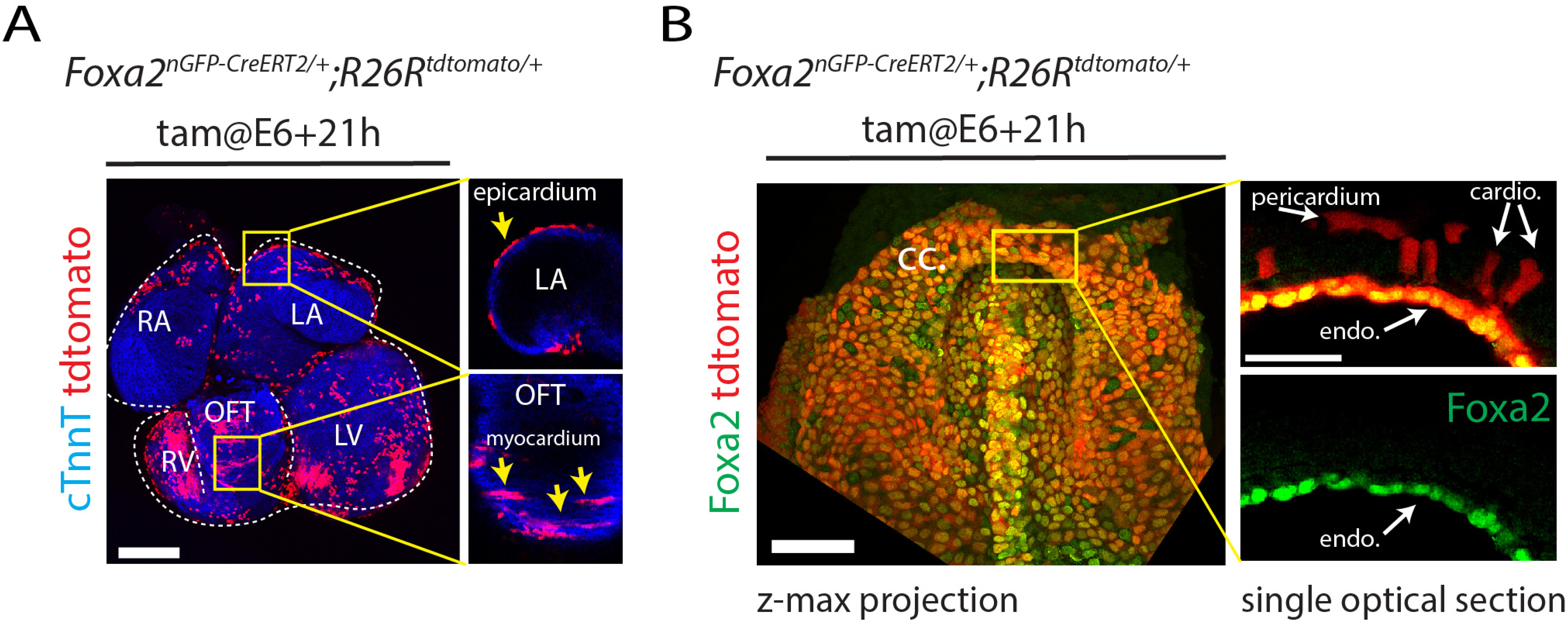
F*o*xa2-expressing cells contribute to the outflow tract myocardium but not to the atria. **(A)** Heart resulting from the administration of tamoxifen at E6+21h. View is ventral. TdTomato-positive cardiomyocytes are absent from the myocardium in the atria, however, contribution to the epicardium (yellow arrow) and myocardium (yellow arrows) in the ventricle and outflow tract is visible. (**B**) E8 embryo resulting from the administration of tamoxifen at E6+21h in *Foxa2^nGPF-CreERT2/+^; R26R^tdTomato/+^* mouse and immunostained for Foxa2 (green). TdTomato-positive cells are localised in the pericardium, cardiomyocytes and endoderm but not in the endocardium. OFT: outflow tract, LA: left atria, RA: right atria, LV: left ventricle, RV: right ventricle, CC: cardiac crescent, cardio: cardiomyocyte, endo: endoderm, Scale bars: 200 µm in (A) and 100 µm in (B).

**S6 Fig.**
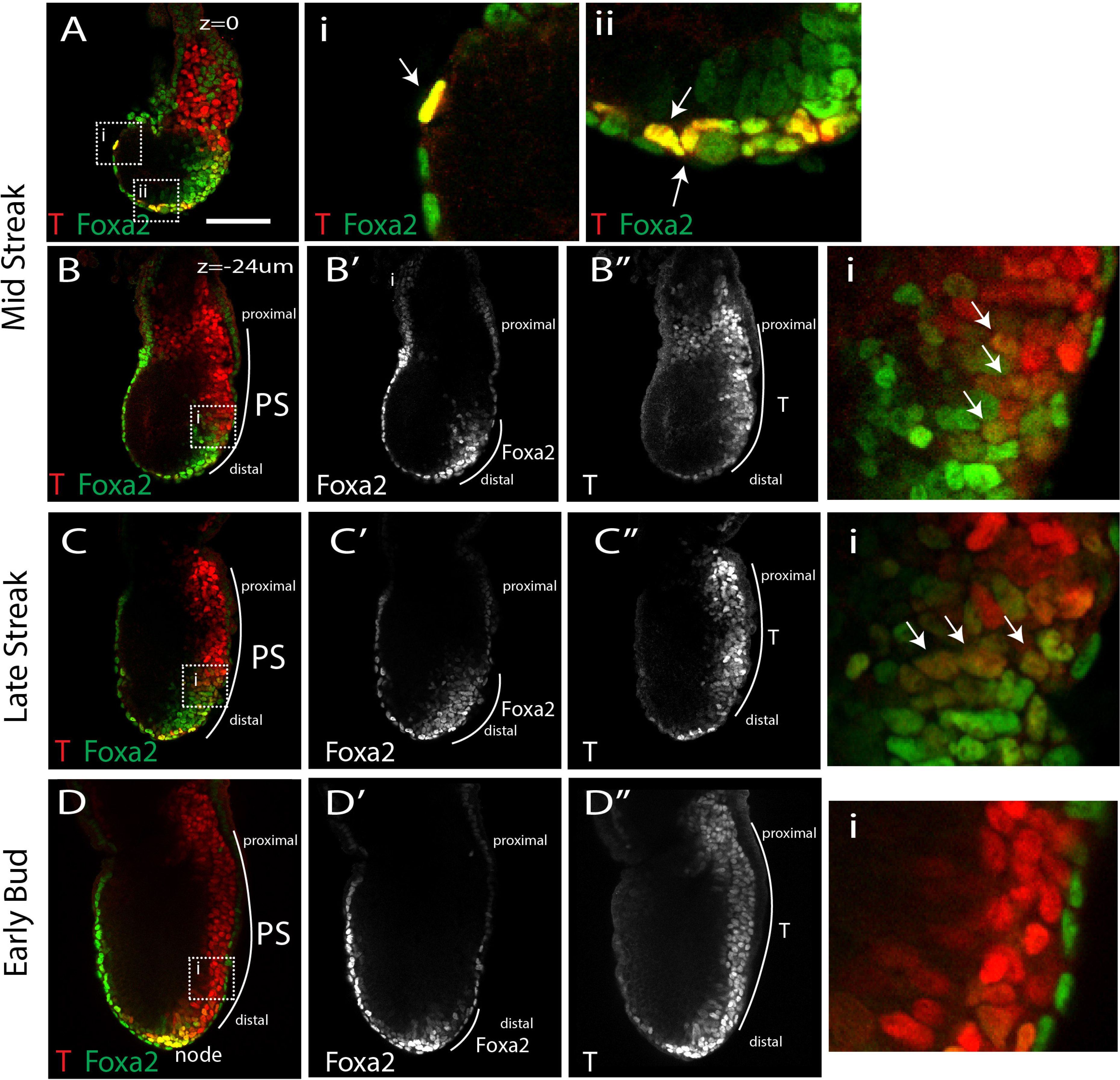
T and Foxa2 colocalise in primitive streak cells. (**A-D**) Single optical sections from same embryos as shown in Fig 4. E6+21h MS (A-A”) and LS (B-B”) and E7+7h “early bud” EB (E-F) embryos are immunostained for T (red) and Foxa2 (green). Views are lateral/slightly posterior. Insets in Ai-ii, Bi and Di show magnified views (A-C). White arrows point to T+/Foxa2+ double positive cells in the definitive endoderm (Ai-ii) at mid-streak position in MS-LS embryos (Bi and Ci). Scale bar: 100 µm.

**S7 Fig.**
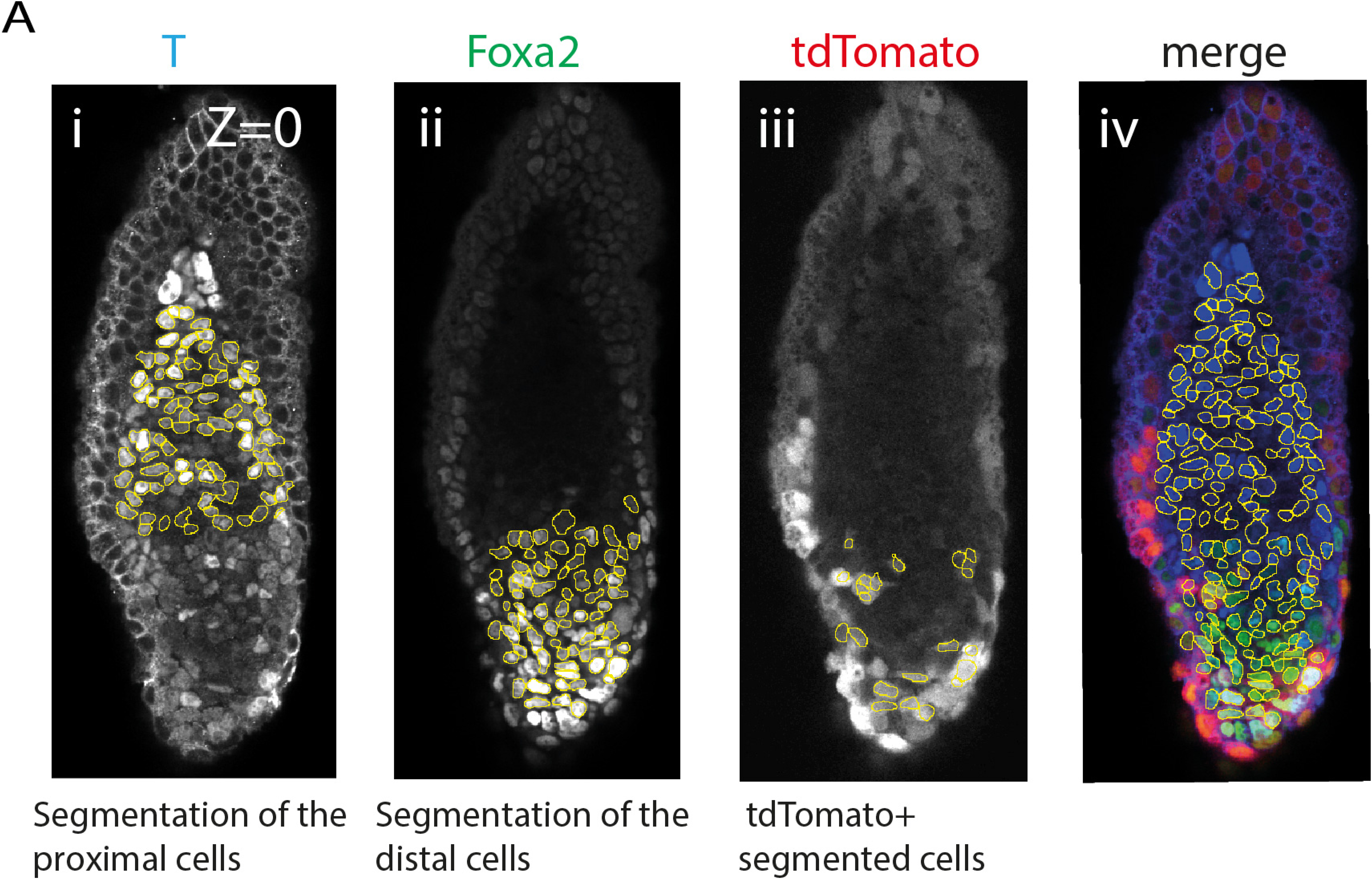
Segmentation of the proximal and distal primitive streak cells. (**A**) Example of a segmented images based on T signal for the proximal cells (i) and Foxa2 signal for the distal cells (ii). Segmentation for only the tdTomato positive cells is shown in (iii). Merge of the two segmented images (i and ii) is shown in (iv).

**S8 Fig.**
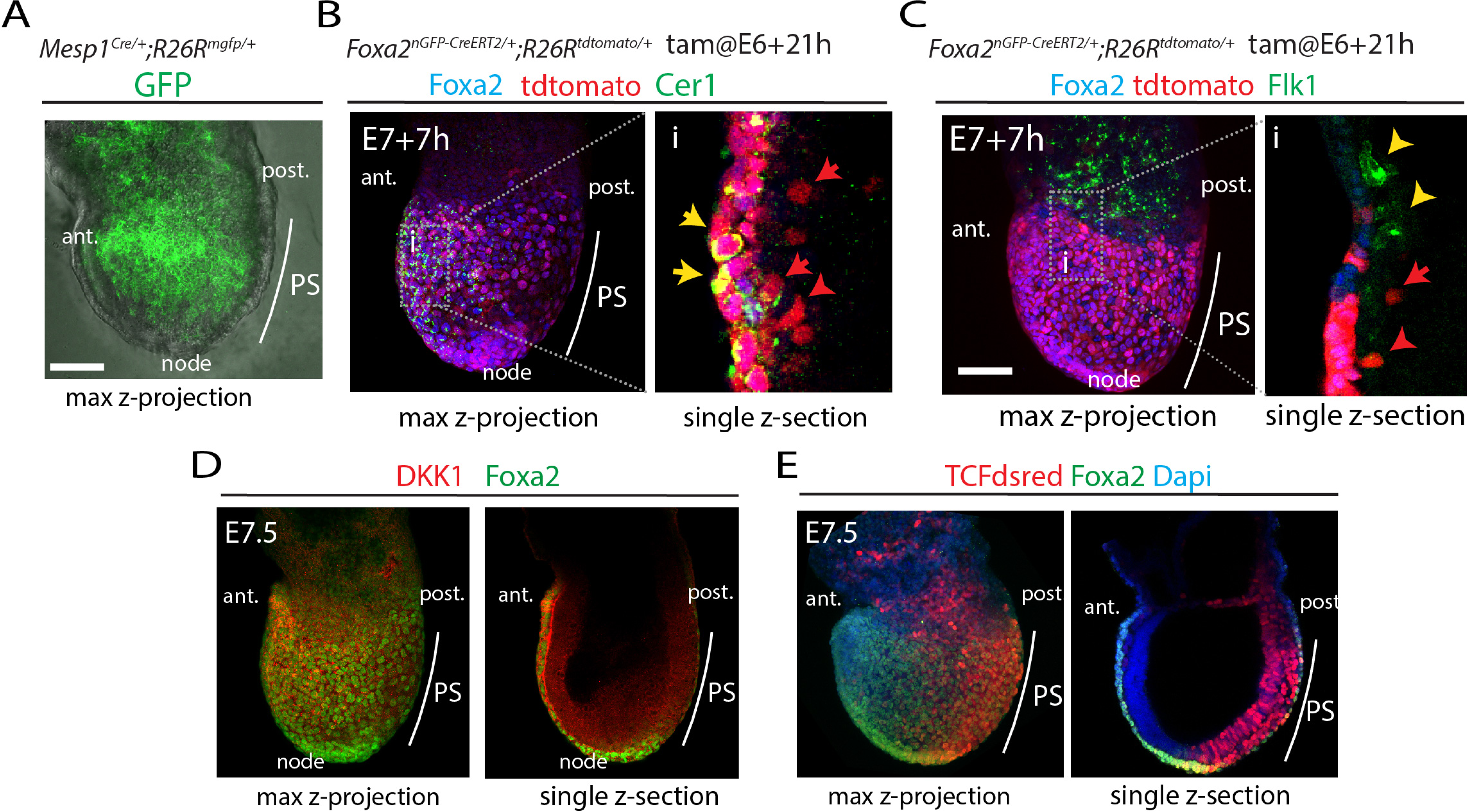
Characterisation of the *Foxa2* lineage positive mesodermal cells. **(A)** Representative *Mesp1^cre/+^;R26R^mGFP/+^* embryo at ∼E7.5. (**B-C**) Representative embryos resulting from the administration of tamoxifen at E6+21h in *Foxa2^nGPF-CreERT2/+^; R26R^tdTomato/+^* immunostained for Foxa2 (bleu) and Cer1 (bleu) (B) or Foxa2 (bleu) and Flk1 (green)(C). Inset in Bi-Ci show magnified view (B-C) in single optical section. (**D**) Representative E7.5 embryo immunostained for DKK1 (red) and Foxa2 (green). (**E**) Representative TCFdsred embryo (red) at E7.5 immunostained for Foxa2 (green). Ant. Anterior, post.: posterior, PS: primitive streak. Scale bar: 100 µm.

**S9 Fig.**
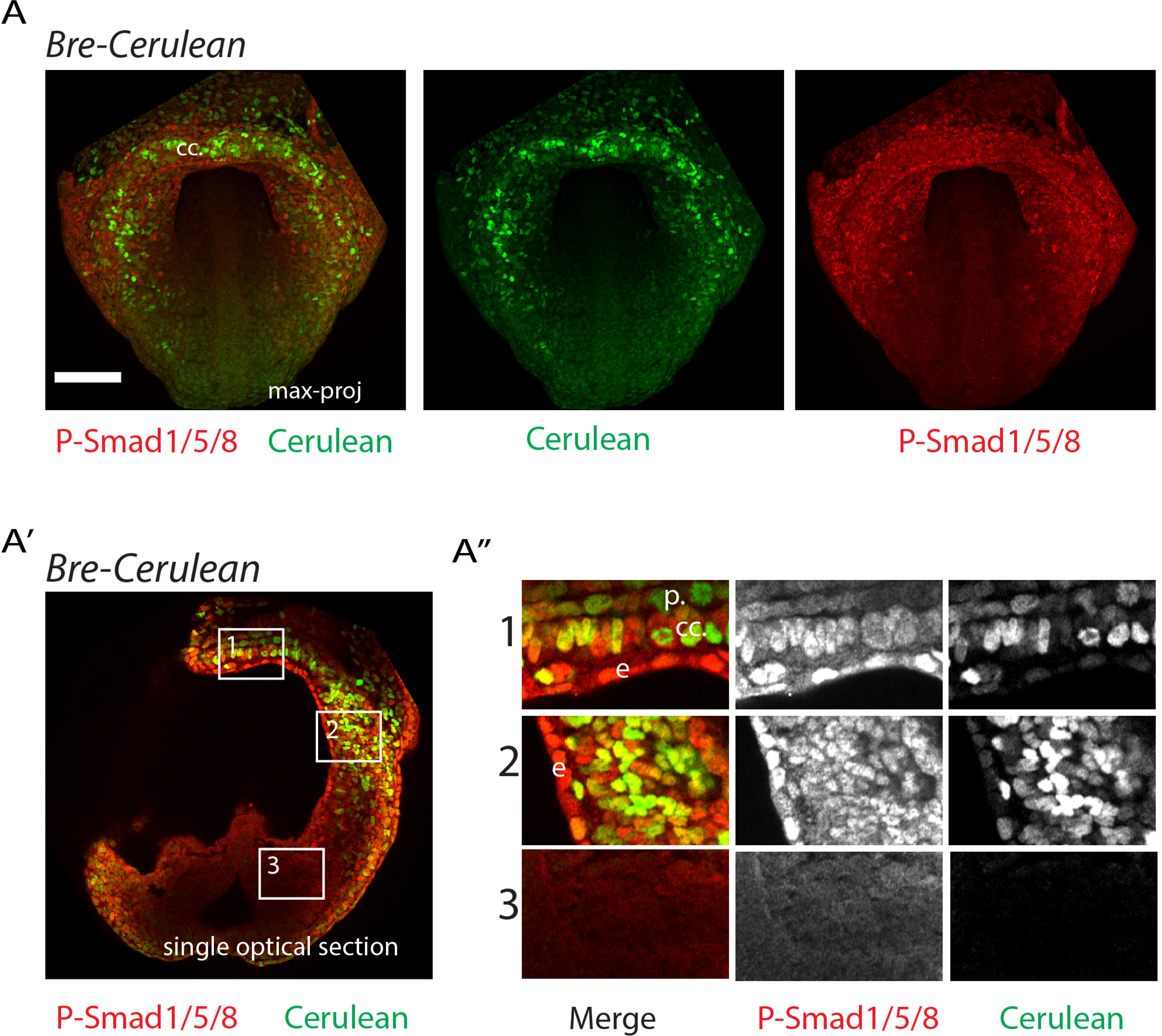
Bre-cerulean line report BMP signalling activity in the mesoderm. (**A-A’’**) Colocalisation of the Cerulean signal and P-Smad1/5/8 in Bre-cerulean embryos at the cardiac crescent stage. (A) z-max proj. (A’) Single optical projection. (A’’) Magnified view form insets in A’. cc.: cardiac crescent. e: endoderm. p: pericardium. Scale bar: 100 µm

**S10 Fig.**
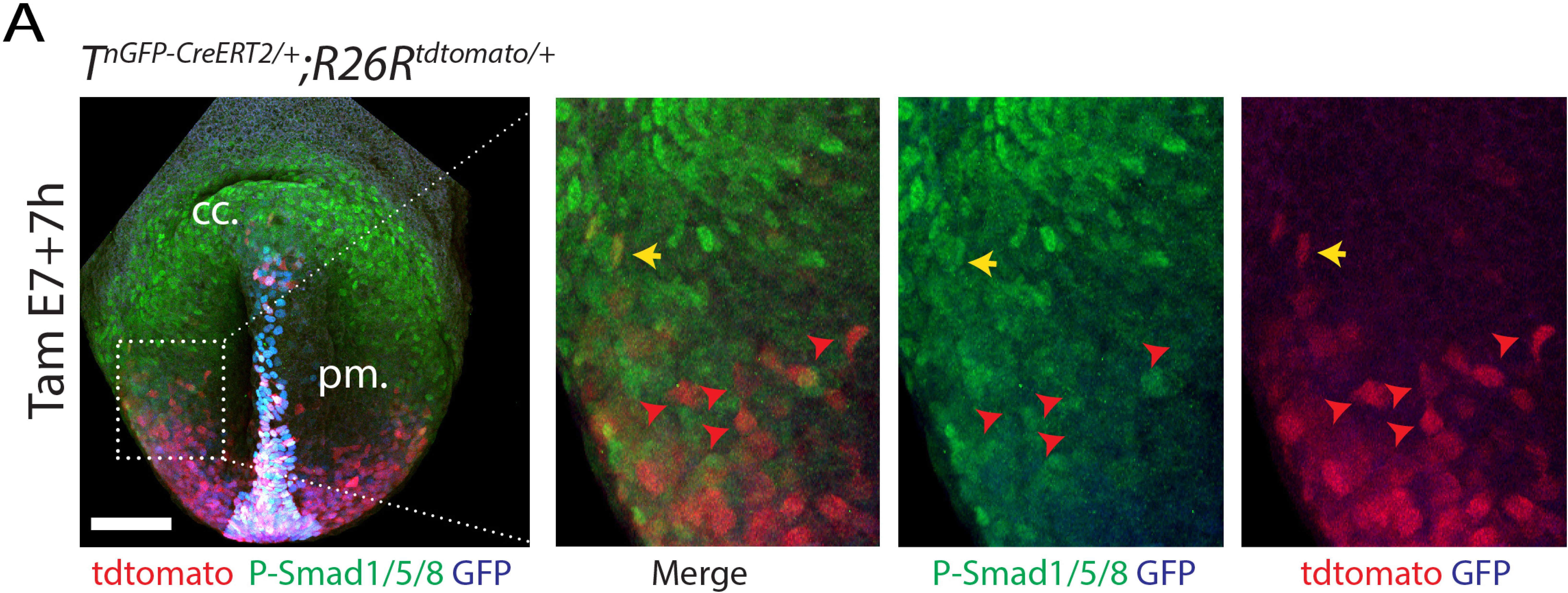
Outflow tract and atrial progenitors are located away from regions with high BMP signalling activity. **(A)** TdTomato localisation in *T^nGPF-CreERT2/+^; R26R^tdTomato/tdTomato^* embryos immunostained against P-Smad1/5/8 following tamoxifen administration at E7+7h. cc: cardiac crescent, pm: pharyngeal mesoderm. Yellow arow points to a Phospho-Smad1/5/8+/tdTomato+ cell, red arrows point to Smad1/5/8-/tdTomato+ cells. Scale bar: 100 µm.

**S11 Fig.**
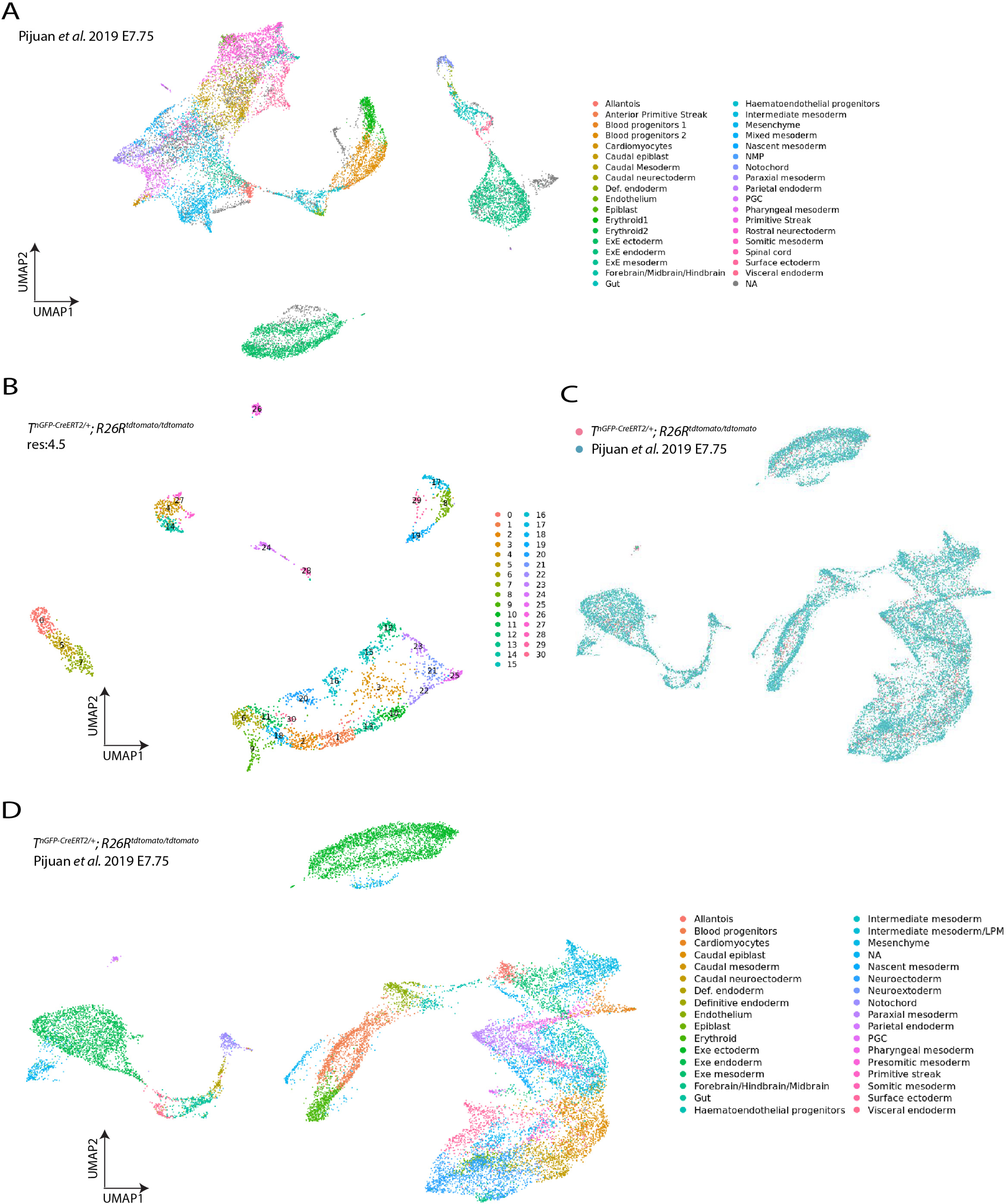
Assignment of cluster identities in scRNA-seq E7+7h dataset. (**A**) UMAP plot of the *Pijuan et al.* E7.75 dataset. (**B**) UMAP plot of the E7+7h *T^nGPF-CreERT2/+^; R26R^tdTomato/tdTomato^* dataset clustered at resolution 4.5. (**C-D**) UMAP plot showing the integrated data from the two scRNA-seq E7+7h *T^nGPF-CreERT2/+^; R26R^tdTomato/tdTomato^* and *Pijuan et al.* E7.75 dataset. Colour codes correspond to the embryonic stage of collection or population identity (**C**) and clusters (**D**). Note, the paraxial mesoderm cluster is split into two sublcusters we named “paraxial mesoderm” and “anterior paraxial mesoderm” based on expression of marker genes (see also Fig 8D, S12A Fig and S7_Source data).

**S12 Fig.**
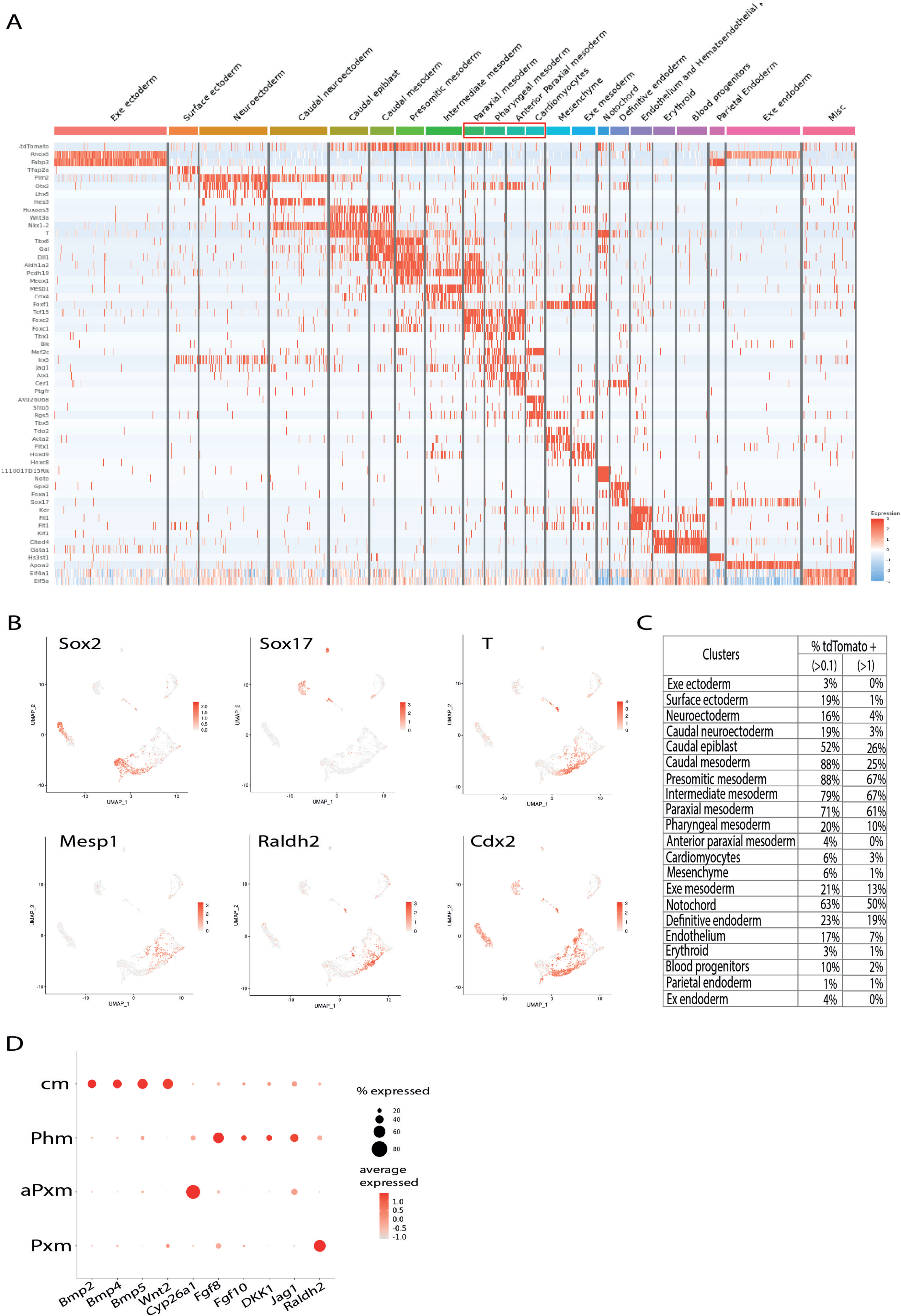
scRNA-seq analysis of the E7+7h *T^nGPF-CreERT2/+^; R26R^tdTomato/tdTomato^* dataset. **(A)** Expression heat map of marker genes (S7_Source data) and tdTomato. Scale indicates z-scored expression values. (**B**) UMAP showing the log normalised counts of selected genes (**C**) Percentage of tdTomato positive cells in each cluster for expression values above 0.1 and 1. (**D**) Dot plot of factors with restricted expression in progenitors. Dot size corresponds to the percentage of cells expressing the feature in each cluster, while the colour represents the average expression level. Cm: cardiomyocytes, Phm: pharyngeal mesoderm, aPxm: anterior paraxial mesoderm, Pxm: paraxial mesoderm.

**S13 Fig.**
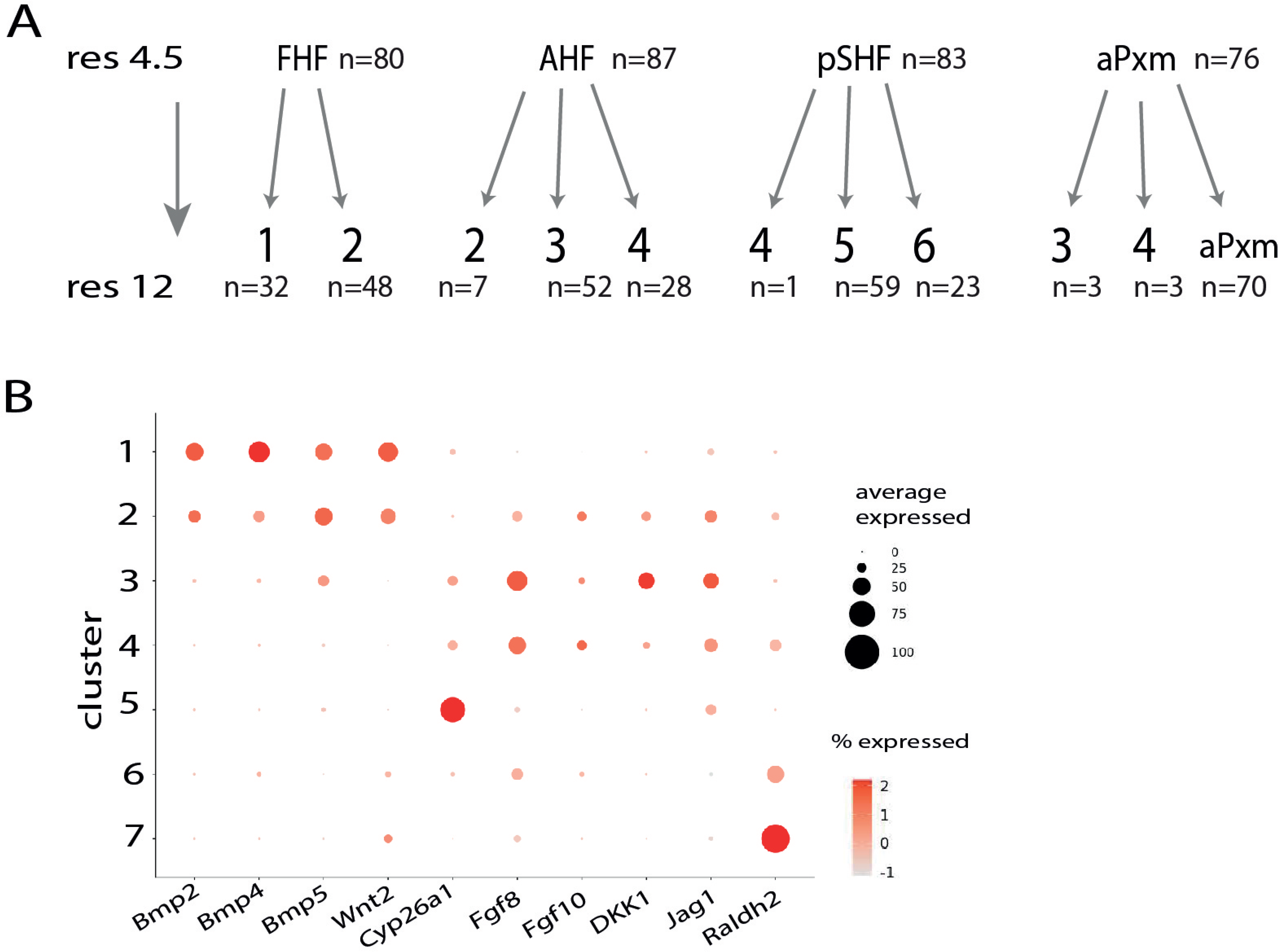
Identification of mesodermal sublclusters and gradient of tdTomato expression. **(A)** Repartition of the cells from the FHF, AHF, pSHF and aPxm cluster to subclucster 1,2,3,4,5 and 6 and aPxm. (**B**) Dot plot of factors with restricted expression in progenitors. Dot size corresponds to the percentage of cells expressing the feature in each cluster, while the colour represents the average expression level. aPxm: anteriorparaxial mesoderm, FHF;first heart field, AHF: anterior heart field, pSHF: posterior second heart field.

**S14 Fig.**
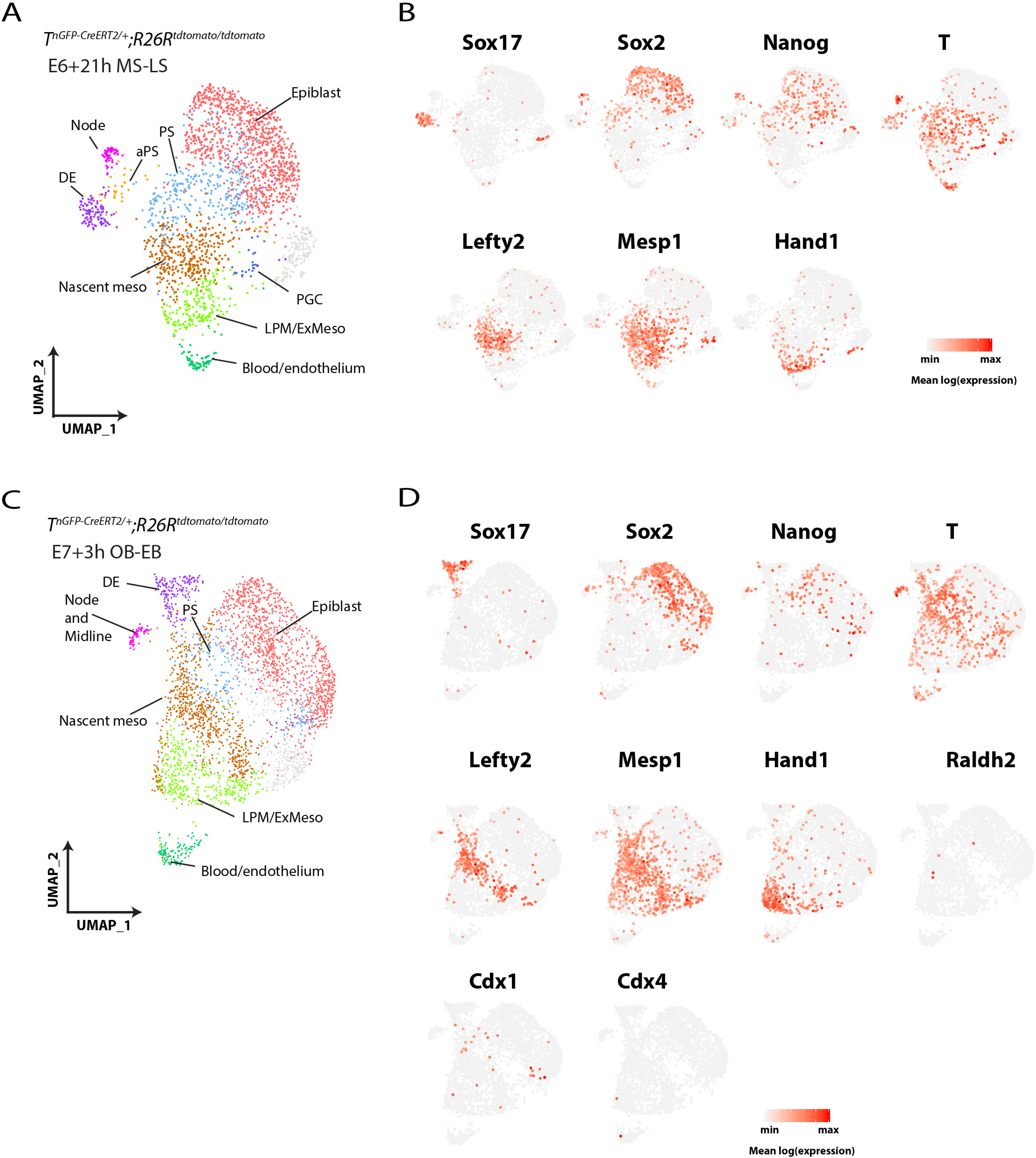
scRNA-seq analysis of the mid-late streak and OB-EB stages. **(A** and **C)** UMAP plot coloured by cluster identity from sc-RNA seq analysis of *T^nGPF-CreERT2/+^; R26R^tdTomato/tdTomato^* embryos at the E6+21h, MS-LS stages (A) and at the E7+3h, OB-EB stages (C). (**B** and **D**) UMAP showing the log normalised counts of selected genes. Colour intensity is proportional to the expression level of a given gene. DE: definitive endoderm, aPS: anterior primitive streak, PS: primitive streak, Nascent meso: nascent mesoderm, LPM/Ex-meso: lateral plate mesoderm and Extra-embryonic mesoderm, mesenchyme. PGC: primordial germs cells. MS: mid-streak, LS: late streak, OB: no bud, EB: early bud.

**S15 Fig.**
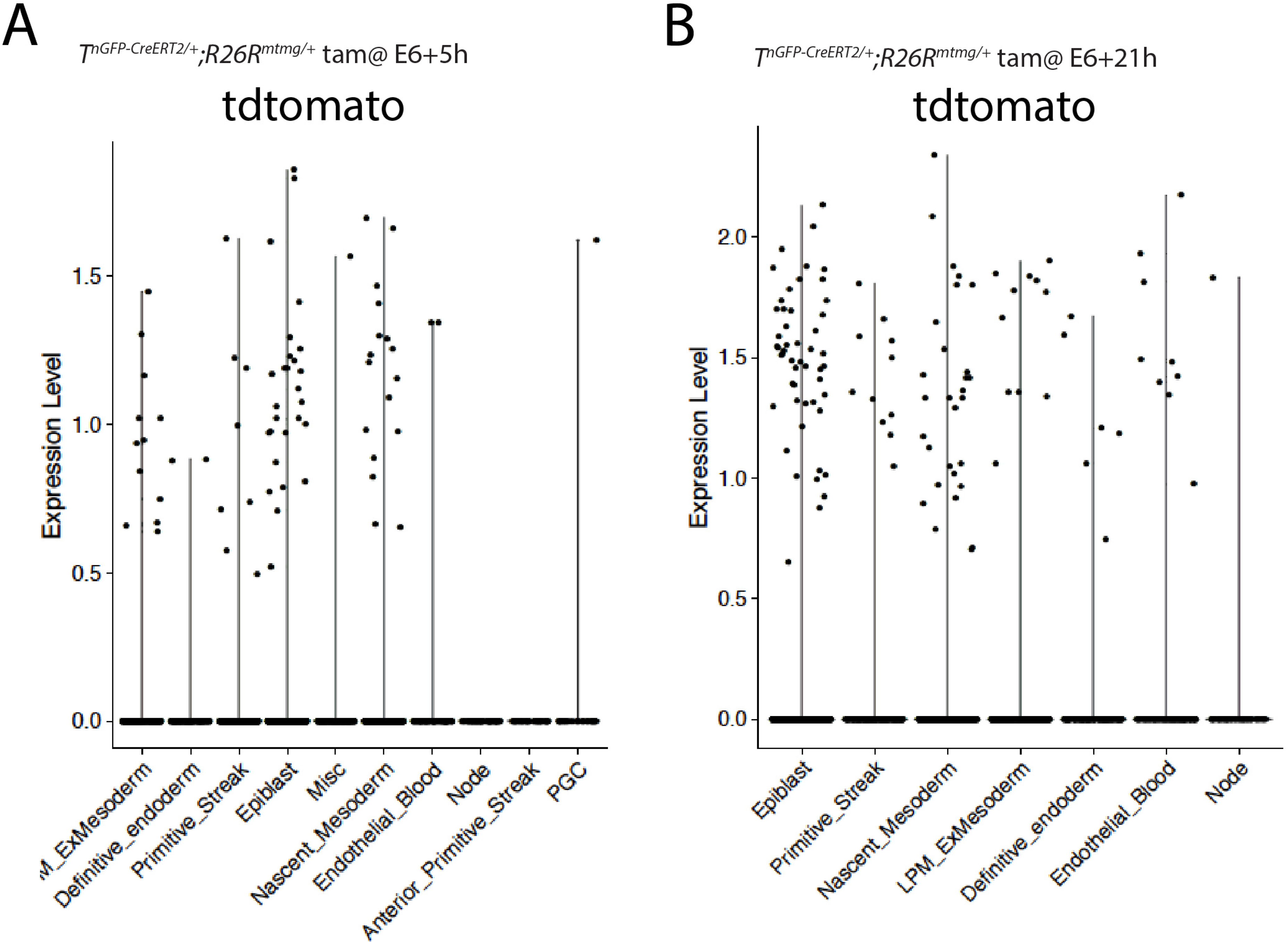
TdTomato reads in *T^nGPF-CreERT2/+^; R26R^tdTomato/tdTomato^* at the mid-late streak and OB-EB stages. **(A-B)** Violin plots showing tdTomato expression for each cluster in *T^nGPF-CreERT2/+^; R26R^tdTomato/tdTomato^* mid-late streak (A) and OB-EB (B) embryos shown in Fig 9A.

**S1_raw image.** Related to Fig 1G

**S2_raw image.** Related to S4 Fig.

**S3_ source data.** Quantification of labelled surface area related to Fig 1E

**S4_source data.** Quantification of T and ERT intensities related to Fig 1K and M

**S5_source data.** Quantification of labelled surface area related to Fig 3D

**S6_source data.** Quantification of T and Foxa2 intensities related to Fig 5E and F.

**S7_source data.** Single cell RNA sequencing data related to Fig 8F and S12 Fig.

**S8_source data.** Quantification of the tdTomato positive cells in each cluster related to Fig 8C and E and S12C Fig.

**S9_source data.** Single cell RNA sequencing data related to Fig 9 and S14 Fig.

**S10_movie.** Related to Fig10.

## Notes

### Competing Interest Statement

The authors have declared no competing interest.

## References

Acampora, D., Annino, A., Puelles, E., Alfano, I., Tuorto, F., and Simeone, A. (2003). OTX1 compensates for OTX2 requirement in regionalisation of anterior neuroectoderm. Gene Expr Patterns 3, 497–501.

Albano, R.M., Arkell, R., Beddington, R.S., and Smith, J.C. (1994). Expression of inhibin subunits and follistatin during postimplantation mouse development: decidual expression of activin and expression of follistatin in primitive streak, somites and hindbrain. Development 120, 803–813.

Alsan, B.H., and Schultheiss, T.M. (2002). Regulation of avian cardiogenesis by Fgf8 signaling. Development 129, 1935–1943.

Andersen, P., Tampakakis, E., Jimenez, D.V., Kannan, S., Miyamoto, M., Shin, H.K., Saberi, A., Murphy, S., Sulistio, E., Chelko, S.P., et al. (2018). Precardiac organoids form two heart fields via Bmp/Wnt signaling. Nat Commun 9, 3140.

Anderson, C., Khan, M.A.F., Wong, F., Solovieva, T., Oliveira, N.M.M., Baldock, R.A., Tickle, C., Burt, D.W., and Stern, C.D. (2016). A strategy to discover new organizers identifies a putative heart organizer. Nat Commun 7, 12656.

Ang, S.L., and Rossant, J. (1994). HNF-3 beta is essential for node and notochord formation in mouse development. Cell 78, 561–574.

Arnold, S.J., and Robertson, E.J. (2009). Making a commitment: cell lineage allocation and axis patterning in the early mouse embryo. Nat Rev Mol Cell Biol 10, 91–103.

Bachiller, D., Klingensmith, J., Kemp, C., Belo, J.A., Anderson, R.M., May, S.R., McMahon, J.A., McMahon, A.P., Harland, R.M., Rossant, J., et al. (2000). The organizer factors Chordin and Noggin are required for mouse forebrain development. Nature 403, 658–661.

Baldini, A., Fulcoli, F.G., and Illingworth, E. (2017). Tbx1: Transcriptional and Developmental Functions. Curr Top Dev Biol 122, 223–243.

Bardot, E., Calderon, D., Santoriello, F., Han, S., Cheung, K., Jadhav, B., Burtscher, I., Artap, S., Jain, R., Epstein, J., et al. (2017). Foxa2 identifies a cardiac progenitor population with ventricular differentiation potential. Nat Commun 8, 14428.

Bertrand, N., Roux, M., Ryckebusch, L., Niederreither, K., Dolle, P., Moon, A., Capecchi, M., and Zaffran, S. (2011). Hox genes define distinct progenitor sub-domains within the second heart field. Dev Biol 353, 266–274.

Blum, M., Gaunt, S.J., Cho, K.W., Steinbeisser, H., Blumberg, B., Bittner, D., and De Robertis, E.M. (1992). Gastrulation in the mouse: the role of the homeobox gene goosecoid. Cell 69, 1097–1106.

Burtscher, I., and Lickert, H. (2009). Foxa2 regulates polarity and epithelialization in the endoderm germ layer of the mouse embryo. Development 136, 1029–1038.

Cai, C.L., Liang, X., Shi, Y., Chu, P.H., Pfaff, S.L., Chen, J., and Evans, S. (2003). Isl1 identifies a cardiac progenitor population that proliferates prior to differentiation and contributes a majority of cells to the heart. Dev Cell 5, 877–889.

Campione, M., Steinbeisser, H., Schweickert, A., Deissler, K., van Bebber, F., Lowe, L.A., Nowotschin, S., Viebahn, C., Haffter, P., Kuehn, M.R., et al. (1999). The homeobox gene Pitx2: mediator of asymmetric left-right signaling in vertebrate heart and gut looping. Development 126, 1225–1234.

Chung, J.H., Whiteley, M., and Felsenfeld, G. (1993). A 5’ element of the chicken beta-globin domain serves as an insulator in human erythroid cells and protects against position effect in Drosophila. Cell 74, 505–514.

Costello, I., Nowotschin, S., Sun, X., Mould, A.W., Hadjantonakis, A.K., Bikoff, E.K., and Robertson, E.J. (2015). Lhx1 functions together with Otx2, Foxa2, and Ldb1 to govern anterior mesendoderm, node, and midline development. Genes Dev 29, 2108–2122.

Costello, I., Pimeisl, I.M., Drager, S., Bikoff, E.K., Robertson, E.J., and Arnold, S.J. (2011). The T-box transcription factor Eomesodermin acts upstream of Mesp1 to specify cardiac mesoderm during mouse gastrulation. Nat Cell Biol 13, 1084–1091.

Crossley, P.H., and Martin, G.R. (1995). The mouse Fgf8 gene encodes a family of polypeptides and is expressed in regions that direct outgrowth and patterning in the developing embryo. Development 121, 439–451.

Cunningham, T.J., Yu, M.S., McKeithan, W.L., Spiering, S., Carrette, F., Huang, C.T., Bushway, P.J., Tierney, M., Albini, S., Giacca, M., et al. (2017). Id genes are essential for early heart formation. Genes Dev 31, 1325–1338.

David, R., Brenner, C., Stieber, J., Schwarz, F., Brunner, S., Vollmer, M., Mentele, E., Muller-Hocker, J., Kitajima, S., Lickert, H., et al. (2008). MesP1 drives vertebrate cardiovascular differentiation through Dkk-1-mediated blockade of Wnt-signalling. Nat Cell Biol 10, 338–345.

Delile, J., Rayon, T., Melchionda, M., Edwards, A., Briscoe, J., and Sagner, A. (2019). Single cell transcriptomics reveals spatial and temporal dynamics of gene expression in the developing mouse spinal cord. Development 146.

de Soysa, T.Y., Ranade, S.S., Okawa, S., Ravichandran, S., Huang, Y., Salunga, H.T., Schricker, A., Del Sol, A., Gifford, C.A., and Srivastava, D. (2019). Single-cell analysis of cardiogenesis reveals basis for organ-level developmental defects. Nature 572, 120–124.

Devine, W.P., Wythe, J.D., George, M., Koshiba-Takeuchi, K., and Bruneau, B.G. (2014). Early patterning and specification of cardiac progenitors in gastrulating mesoderm. Elife 2014;3:e03848

Ding, J., Yang, L., Yan, Y.T., Chen, A., Desai, N., Wynshaw-Boris, A., and Shen, M.M. (1998). Cripto is required for correct orientation of the anterior-posterior axis in the mouse embryo. Nature 395, 702–707.

Doe, C.Q. (2017). Temporal Patterning in the Drosophila CNS. Annu Rev Cell Dev Biol 33, 219–240.

Dolle, P., Fraulob, V., Gallego-Llamas, J., Vermot, J., and Niederreither, K. (2010). Fate of retinoic acid-activated embryonic cell lineages. Dev Dyn 239, 3260–3274.

Dono, R., Scalera, L., Pacifico, F., Acampora, D., Persico, M.G., and Simeone, A. (1993). The murine cripto gene: expression during mesoderm induction and early heart morphogenesis. Development 118, 1157–1168.

Dorey, K., and Amaya, E. (2010). FGF signalling: diverse roles during early vertebrate embryogenesis. Development 137, 3731–3742.

Downs, K.M., and Davies, T. (1993). Staging of gastrulating mouse embryos by morphological landmarks in the dissecting microscope. Development 118, 1255–1266.

Evan S. Bardot, Bharati Jadhav, Nadeera Wickramasinghe, Amélie Rezza, Michael Rendl, Andrew J. Sharp, Nicole C. Dubois. Notch Signalling Commits Mesoderm to the Cardiac Lineage. bioRxiv 2020.02.20.958348; doi: https://doi.org/10.1101/2020.02.20.958348

Filipe, M., Goncalves, L., Bento, M., Silva, A.C., and Belo, J.A. (2006). Comparative expression of mouse and chicken Shisa homologues during early development. Dev Dyn 235, 2567–2573.

Frank, D.U., Elliott, S.A., Park, E.J., Hammond, J., Saijoh, Y., and Moon, A.M. (2007). System for inducible expression of cre-recombinase from the Foxa2 locus in endoderm, notochord, and floor plate. Dev Dyn 236, 1085–1092.

Garcia-Martinez, V., and Schoenwolf, G.C. (1993). Primitive-streak origin of the cardiovascular system in avian embryos. Dev Biol 159, 706–719.

Guo, Y., Dorn, T., Kuhl, S.J., Linnemann, A., Rothe, M., Pfister, A.S., Vainio, S., Laugwitz, K.L., Moretti, A., and Kuhl, M. (2019). The Wnt inhibitor Dkk1 is required for maintaining the normal cardiac differentiation program in Xenopus laevis. Dev Biol 449, 1–13.

Guzzetta, A., Koska, M., Rowton, M., Sullivan, K.R., Jacobs-Li, J., Kweon, J., Hidalgo, H., Eckart, H., Hoffmann, A.D., Back, R., et al. (2020). Hedgehog-FGF signaling axis patterns anterior mesoderm during gastrulation. Proc Natl Acad Sci U S A.

Hoang, B.H., Thomas, J.T., Abdul-Karim, F.W., Correia, K.M., Conlon, R.A., Luyten, F.P., and Ballock, R.T. (1998). Expression pattern of two Frizzled-related genes, Frzb-1 and Sfrp-1, during mouse embryogenesis suggests a role for modulating action of Wnt family members. Dev Dyn 212, 364–372.

Hochgreb, T., Linhares, V.L., Menezes, D.C., Sampaio, A.C., Yan, C.Y., Cardoso, W.V., Rosenthal, N., and Xavier-Neto, J. (2003). A caudorostral wave of RALDH2 conveys anteroposterior information to the cardiac field. Development 130, 5363–5374.

Huynh, T., Chen, L., Terrell, P., and Baldini, A. (2007). A fate map of Tbx1 expressing cells reveals heterogeneity in the second cardiac field. Genesis 45, 470–475.

Ilagan, R., Abu-Issa, R., Brown, D., Yang, Y.P., Jiao, K., Schwartz, R.J., Klingensmith, J., and Meyers, E.N. (2006). Fgf8 is required for anterior heart field development. Development 133, 2435–2445.

Imuta, Y., Kiyonari, H., Jang, C.W., Behringer, R.R., and Sasaki, H. (2013). Generation of knock-in mice that express nuclear enhanced green fluorescent protein and tamoxifen-inducible Cre recombinase in the notochord from Foxa2 and T loci. Genesis 51, 210–218.

Ivanovitch, K., Temino, S., and Torres, M. (2017). Live imaging of heart tube development in mouse reveals alternating phases of cardiac differentiation and morphogenesis. Elife 6. eLife 2017;6:e30668

Keegan, B.R., Meyer, D., and Yelon, D. (2004). Organization of cardiac chamber progenitors in the zebrafish blastula. Development 131, 3081–3091.

Kelly, R.G., Brown, N.A., and Buckingham, M.E. (2001). The arterial pole of the mouse heart forms from Fgf10-expressing cells in pharyngeal mesoderm. Dev Cell 1, 435–440.

Kinder, S.J., Tsang, T.E., Quinlan, G.A., Hadjantonakis, A.K., Nagy, A., and Tam, P.P. (1999). The orderly allocation of mesodermal cells to the extraembryonic structures and the anteroposterior axis during gastrulation of the mouse embryo. Development 126, 4691–4701.

Kinder, S.J., Tsang, T.E., Wakamiya, M., Sasaki, H., Behringer, R.R., Nagy, A., and Tam, P.P. (2001). The organizer of the mouse gastrula is composed of a dynamic population of progenitor cells for the axial mesoderm. Development 128, 3623–3634.

Klaus, A., Saga, Y., Taketo, M.M., Tzahor, E., and Birchmeier, W. (2007). Distinct roles of Wnt/beta-catenin and Bmp signaling during early cardiogenesis. Proc Natl Acad Sci U S A 104, 18531–18536.

Lee, C.S., and Fan, C.M. (2001). Embryonic expression patterns of the mouse and chick Gas1 genes. Mech Dev 101, 293–297.

Lee, J.H., Protze, S.I., Laksman, Z., Backx, P.H., and Keller, G.M. (2017). Human Pluripotent Stem Cell-Derived Atrial and Ventricular Cardiomyocytes Develop from Distinct Mesoderm Populations. Cell Stem Cell 21, 179–194 e174.

Lescroart, F., Chabab, S., Lin, X., Rulands, S., Paulissen, C., Rodolosse, A., Auer, H., Achouri, Y., Dubois, C., Bondue, A., et al. (2014). Early lineage restriction in temporally distinct populations of Mesp1 progenitors during mammalian heart development. Nat Cell Biol 16, 829–840.

Lescroart, F., Wang, X., Lin, X., Swedlund, B., Gargouri, S., Sanchez-Danes, A., Moignard, V., Dubois, C., Paulissen, C., Kinston, S., et al. (2018). Defining the earliest step of cardiovascular lineage segregation by single-cell RNA-seq. Science 359, 1177–1181.

Lough, J., and Sugi, Y. (2000). Endoderm and heart development. Dev Dyn 217, 327–342.

Madisen, L., Zwingman, T.A., Sunkin, S.M., Oh, S.W., Zariwala, H.A., Gu, H., Ng, L.L., Palmiter, R.D., Hawrylycz, M.J., Jones, A.R., et al. (2010). A robust and high-throughput Cre reporting and characterization system for the whole mouse brain. Nat Neurosci 13, 133–140.

Meilhac, S.M., and Buckingham, M.E. (2018). The deployment of cell lineages that form the mammalian heart. Nat Rev Cardiol 15, 705–724.

Meilhac, S.M., Esner, M., Kelly, R.G., Nicolas, J.F., and Buckingham, M.E. (2004). The clonal origin of myocardial cells in different regions of the embryonic mouse heart. Dev Cell 6, 685–698.

Mendjan, S., Mascetti, V.L., Ortmann, D., Ortiz, M., Karjosukarso, D.W., Ng, Y., Moreau, T., and Pedersen, R.A. (2014). NANOG and CDX2 pattern distinct subtypes of human mesoderm during exit from pluripotency. Cell Stem Cell 15, 310–325.

Metzis, V., Steinhauser, S., Pakanavicius, E., Gouti, M., Stamataki, D., Ivanovitch, K., Watson, T., Rayon, T., Mousavy Gharavy, S.N., Lovell-Badge, R., et al. (2018). Nervous System Regionalization Entails Axial Allocation before Neural Differentiation. Cell 175, 1105–1118 e1117.

Monkley, S.J., Delaney, S.J., Pennisi, D.J., Christiansen, J.H., and Wainwright, B.J. (1996). Targeted disruption of the Wnt2 gene results in placentation defects. Development 122, 3343–3353.

Morgani, S.M., Metzger, J.J., Nichols, J., Siggia, E.D., and Hadjantonakis, A.K. (2018). Micropattern differentiation of mouse pluripotent stem cells recapitulates embryo regionalized cell fate patterning. Elife 7.

Muzumdar, M.D., Tasic, B., Miyamichi, K., Li, L., and Luo, L. (2007). A global double-fluorescent Cre reporter mouse. Genesis 45, 593–605.

Nam, J.S., Turcotte, T.J., and Yoon, J.K. (2007). Dynamic expression of R-spondin family genes in mouse development. Gene Expr Patterns 7, 306–312.

Nandkishore, N., Vyas, B., Javali, A., Ghosh, S., and Sambasivan, R. (2018). Divergent early mesoderm specification underlies distinct head and trunk muscle programmes in vertebrates. Development 145.

Phillips, M.D., Mukhopadhyay, M., Poscablo, C., and Westphal, H. (2011). Dkk1 and Dkk2 regulate epicardial specification during mouse heart development. Int J Cardiol 150, 186–192.

Pijuan-Sala, B., Griffiths, J.A., Guibentif, C., Hiscock, T.W., Jawaid, W., Calero-Nieto, F.J., Mulas, C., Ibarra-Soria, X., Tyser, R.C.V., Ho, D.L.L., et al. (2019). A single-cell molecular map of mouse gastrulation and early organogenesis. Nature 566, 490–495.

Pimeisl, I.M., Tanriver, Y., Daza, R.A., Vauti, F., Hevner, R.F., Arnold, H.H., and Arnold, S.J. (2013). Generation and characterization of a tamoxifen-inducible Eomes(CreER) mouse line. Genesis 51, 725–733.

Przemeck, G.K., Heinzmann, U., Beckers, J., and Hrabe de Angelis, M. (2003). Node and midline defects are associated with left-right development in Delta1 mutant embryos. Development 130, 3–13.

Rivera-Perez, J.A., and Magnuson, T. (2005). Primitive streak formation in mice is preceded by localized activation of Brachyury and Wnt3. Dev Biol 288, 363–371.

Roux, M., Laforest, B., Capecchi, M., Bertrand, N., and Zaffran, S. (2015). Hoxb1 regulates proliferation and differentiation of second heart field progenitors in pharyngeal mesoderm and genetically interacts with Hoxa1 during cardiac outflow tract development. Dev Biol 406, 247–258.

Santos-Ledo, A., Washer, S., Dhanaseelan, T., Eley, L., Alqatani, A., Chrystal, P.W., Papoutsi, T., Henderson, D.J., and Chaudhry, B. (2020). Alternative splicing of jnk1a in zebrafish determines first heart field ventricular cardiomyocyte numbers through modulation of hand2 expression. PLoS Genet 16, e1008782.

Schindelin, J., Arganda-Carreras, I., Frise, E., Kaynig, V., Longair, M., Pietzsch, T., Preibisch, S., Rueden, C., Saalfeld, S., Schmid, B., et al. (2012). Fiji: an open-source platform for biological-image analysis. Nat Methods 9, 676–682.

Schultheiss, T.M., Burch, J.B., and Lassar, A.B. (1997). A role for bone morphogenetic proteins in the induction of cardiac myogenesis. Genes Dev 11, 451–462.

Shimono, A., and Behringer, R.R. (1999). Isolation of novel cDNAs by subtractions between the anterior mesendoderm of single mouse gastrula stage embryos. Dev Biol 209, 369–380.

Sirbu, I.O., Zhao, X., and Duester, G. (2008). Retinoic acid controls heart anteroposterior patterning by down-regulating Isl1 through the Fgf8 pathway. Dev Dyn 237, 1627–1635.

Spater, D., Abramczuk, M.K., Buac, K., Zangi, L., Stachel, M.W., Clarke, J., Sahara, M., Ludwig, A., and Chien, K.R. (2013). A HCN4+ cardiomyogenic progenitor derived from the first heart field and human pluripotent stem cells. Nat Cell Biol 15, 1098–1106.

Stuart, T., Butler, A., Hoffman, P., Hafemeister, C., Papalexi, E., Mauck, W.M., 3rd, Hao, Y., Stoeckius, M., Smibert, P., and Satija, R. (2019). Comprehensive Integration of Single-Cell Data. Cell 177, 1888–1902 e1821.

Thomas, P.Q., Brown, A., and Beddington, R.S. (1998). Hex: a homeobox gene revealing peri-implantation asymmetry in the mouse embryo and an early transient marker of endothelial cell precursors. Development 125, 85–94.

Truett, G.E., Heeger, P., Mynatt, R.L., Truett, A.A., Walker, J.A., and Warman, M.L. (2000). Preparation of PCR-quality mouse genomic DNA with hot sodium hydroxide and tris (HotSHOT). Biotechniques 29, 52, 54.

Tuazon, F.B., and Mullins, M.C. (2015). Temporally coordinated signals progressively pattern the anteroposterior and dorsoventral body axes. Semin Cell Dev Biol 42, 118–133.

Tyser, R.C.V., Ibarra-Soria, X., McDole, K., S, A.J., Godwin, J., van den Brand, T.A.H., Miranda, A.M.A., Scialdone, A., Keller, P.J., Marioni, J.C., et al. (2021). Characterization of a common progenitor pool of the epicardium and myocardium. Science.

Wianny, F., Real, F.X., Mummery, C.L., Van Rooijen, M., Lahti, J., Samarut, J., and Savatier, P. (1998). G1-phase regulators, cyclin D1, cyclin D2, and cyclin D3: up-regulation at gastrulation and dynamic expression during neurulation. Dev Dyn 212, 49–62.

Williams, R., Lendahl, U., and Lardelli, M. (1995). Complementary and combinatorial patterns of Notch gene family expression during early mouse development. Mech Dev 53, 357–368.

Xavier-Neto, J., Neville, C.M., Shapiro, M.D., Houghton, L., Wang, G.F., Nikovits, W., Jr., Stockdale, F.E., and Rosenthal, N. (1999). A retinoic acid-inducible transgenic marker of sino-atrial development in the mouse heart. Development 126, 2677–2687.

Yutzey, K.E., and Bader, D. (1995). Diversification of cardiomyogenic cell lineages during early heart development. Circ Res 77, 216–219.

Zhang Q, Carlin D, Zhu F, Cattaneo P, Ideker T, Evans SM., Bloomekatz J, Chi NC. (2021) Unveiling unexpected complexity and multipotentiality of early heart fields bioRxiv 2021.01.08.425950; doi: https://doi.org/10.1101/2021.01.08.425950

Zhao, Q., Behringer, R.R., and de Crombrugghe, B. (1996). Prenatal folic acid treatment suppresses acrania and meroanencephaly in mice mutant for the Cart1 homeobox gene. Nat Genet 13, 275–283.

