## Supplementary figures and images for "Ventricular, atrial and outflow tract heart progenitors arise from spatially and molecularly distinct regions of the primitive streak"

### S1_raw image

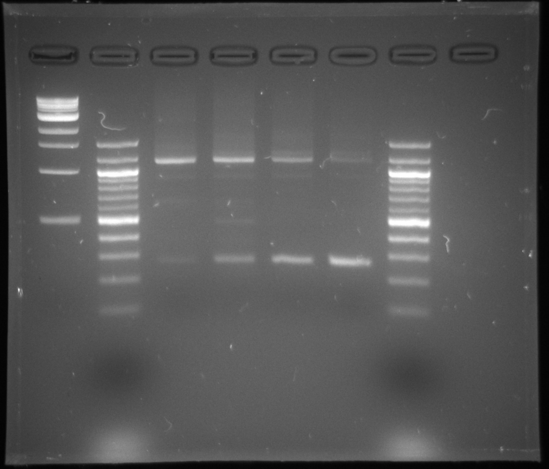

### S2_raw image

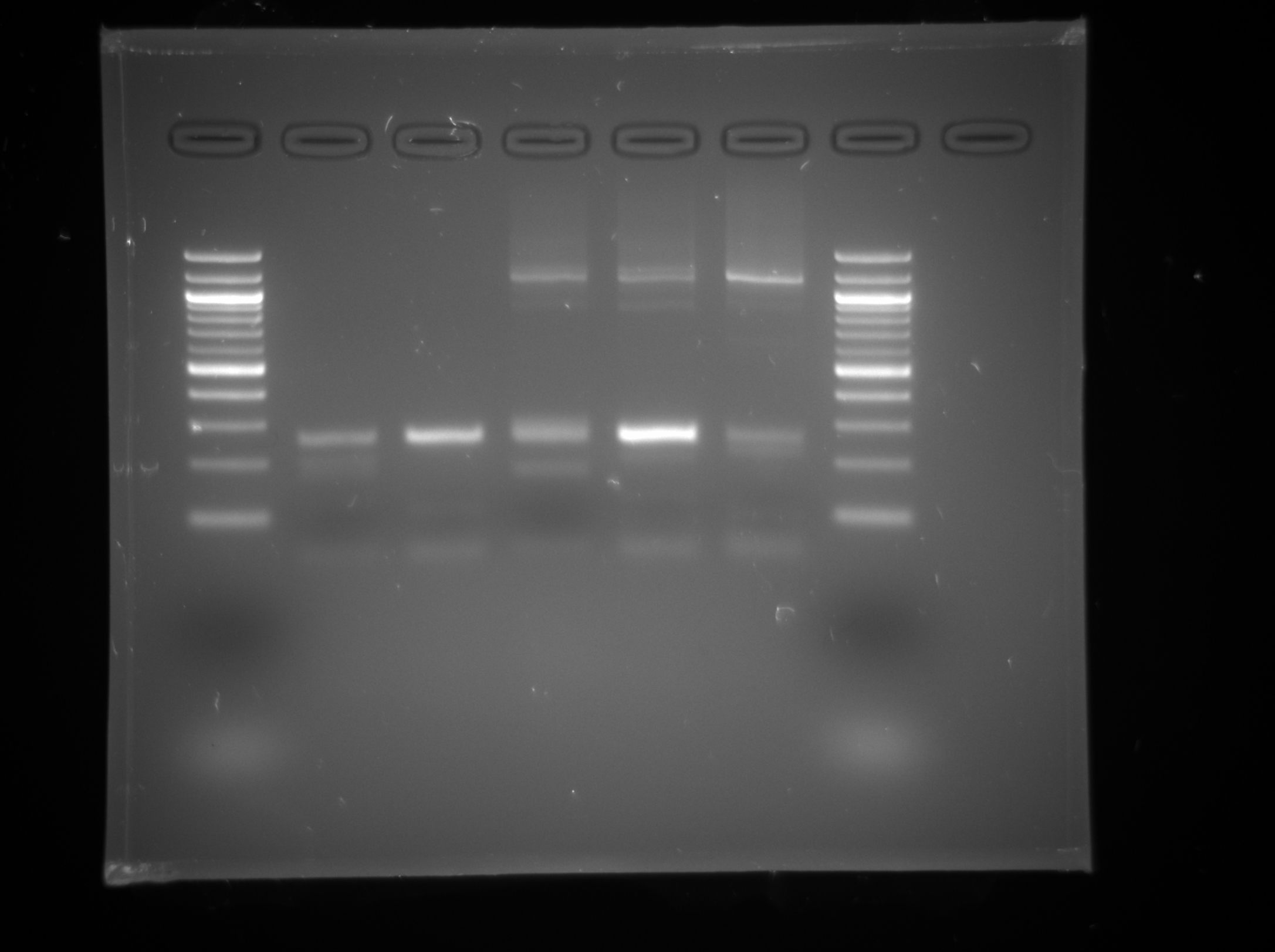
